# A comprehensive analysis of Transcribed Ultra Conserved Regions uncovers important regulatory functions of novel non-coding transcripts in gliomas

**DOI:** 10.1101/2023.09.12.557444

**Authors:** Myron K Gibert, Ying Zhang, Shekhar Saha, Guruprasad Varma Konduru, Pawel Marcinkiewicz, Collin Dube, Kadie Hudson, Yunan Sun, Sylwia Bednarek, Bilhan Chagari, Aditya Sarkar, Christian Roig-Laboy, Natalie Neace, Karim Saoud, Initha Setiady, Farina Hanif, David Schiff, Benjamin Kefas, Markus Hafner, Pankaj Kumar, Roger Abounader

**Affiliations:** University of Virginia Departments of Microbiology, Immunology & Cancer Biology, Charlottesville, VA, 22908, USA; Neurology, Charlottesville, VA, 22908, USA; Cancer Center, Charlottesville, VA, 22908, USA; Department of Public Health Sciences and Bioinformatics Core, Charlottesville, VA, 22908, USA; Department of Biochemistry, Dow International Medical College, Dow University of Health Sciences, Karachi, 75270, Pakistan; National Institutes of Health, Bethesda, MD, USA

## Abstract

Transcribed Ultra-Conserved Regions (TUCRs) represent a severely understudied class of putative non-coding RNAs (ncRNAs) that are 100% conserved across multiple species. We performed the first-ever analysis of TUCRs in glioblastoma (GBM) and low-grade gliomas (LGG). We leveraged large human datasets to identify the genomic locations, chromatin accessibility, transcription, differential expression, correlation with survival, and predicted functions of all 481 TUCRs, and identified TUCRs that are relevant to glioma biology. Of these, we investigated the expression, function, and mechanism of action of the most highly upregulated intergenic TUCR, uc.110, identifying it as a new tumor enhancer. Uc.110 was highly overexpressed in GBM and LGG, where it promoted malignancy and tumor growth. Uc.110 activated the WNT pathway by upregulating the expression of membrane frizzled-related protein (MFRP), by sponging the tumor suppressor microRNA miR-544. This pioneering study shows important roles for TUCRs in gliomas and provides an extensive database and novel methods for future TUCR research.

## INTRODUCTION

Transcribed Ultra-conserved Regions (TUCRs) represent 481 unique transcribed RNA molecules that are “ultraconserved” across multiple species, including human, mouse (100%), rat (100%), dog (98%), and chicken (95%) genomes. [1] TUCR expression has been found to be highly deregulated in some cancers. Because of their ultra-conservation and their deregulation, it is believed that TUCRs may have important regulatory roles in cancer. [2–11] About ∼90% of the genome is transcribed, but only ∼2 percent of the transcriptome is translated. [2] The remainder of the transcriptome is made up of non-coding elements that serve key regulatory roles. Of these elements, long non-coding RNAs (lncRNAs) serve as important regulators of malignancy and potential therapeutic targets in cancer. [2, 12–19] Due to their size and lack of known associated protein products, it has been suggested that many TUCRs, especially the intergenic ones, may function as lncRNAs, while those within introns or spanning introns and exons of protein coding genes may represent either lncRNAs or ultraconserved regions of the coding gene without being independent transcripts themselves. For example, many homeobox (HOX) genes contain ultraconserved regions.[2] Yet, the putative existence of “ultraconserved” lncRNAs is particularly significant, as lncRNAs are typically poorly conserved as a class of molecules.[2] Very little is known about TUCRs. [2] In particular, the literature elucidating the expressions, functions, and mechanisms of action of TUCRs in glioblastoma (GBM) and low-grade glioma (LGG) is nonexistent. To address this gap in knowledge, we leveraged large human glioma datasets to identify the genomic locations, chromatin accessibility, transcription, differential expression, correlation with survival, and predicted functions of all 481 TUCRs, and identified TUCRs that are relevant to glioma biology (Supplementary Figure 1A). GBM and LGG represent over 80% of primary malignant brain tumors in humans, of which GBM is the deadliest, with a median survival of approximately 15 months. [20–28]

We performed expression, survival, chromatin accessibility, and functional predictions, for all 481 TUCRs. Furthermore, we prioritized a further in depth investigation of the expression, function, and mechanism of action, of the most highly upregulated intergenic TUCR, uc.110, identifying it as a new tumor promoter, utilizing both novel and established [2] computational and experimental approaches. Uc.110 was highly overexpressed in GBM and LGG, where it promoted malignancy parameters and tumor growth. Uc.110 activated the WNT pathway by upregulating the expression of membrane frizzled-related protein (MFRP), by sponging the tumor suppressor microRNA miR-544. This work shows important roles for TUCRs in gliomas and provides an extensive database and novel methods for future TUCR research in any disease context.

## RESULTS

### TUCRs are encoded throughout the genome, resistant to variation, and actively transcribed

We analyzed TUCR genomic locations published in Bejerano et al, [1] using hg38 genome coordinates lifted over from the provided hg19 coordinates. We found that some TUCRs are exonic and are contained within an exon of the “host” gene. Others are contained within an intron instead. Some TUCRs straddle a region that spans exonic and intronic regions of the host gene (exonic/intronic), and others are not contained within a known genetic element at all (intergenic) (Supplementary Figure 1B). We manually annotated each TUCR using a combination of UCSC Genome Browser tracks, [29, 30] Quinlan Laboratory’s bedtools, [31, 32] and TUCR genomic locations lifted over to hg38 from hg19. [1] We therefore updated the initial annotations by Bejerano et al, 2004 [1] and subsequently updated annotations by Calin et al, 2007 [4] to now include 45 exonic, 231 intronic, 68 intronic/exonic, and 137 intergenic TUCRs (Supplementary Figure 1C). Intragenic TUCRs (exonic, intronic, and exonic/intronic) tend to congregate in protein coding genes, while intergenic TUCRs likely represent independent and potentially novel transcripts. (Supplementary Figure 1D). There were no annotated TUCRs on chromosome 21 (chr21), the Y chromosome (chrY) or in the mitochondrial DNA (chrM) as previously described by Bejerano et al [1] and Calin et al [4] (Supplementary Figure 1E). Detailed TUCR annotation information for all 481 TUCRs is provided in the supplementary materials (Supplementary Table 1).

We also investigated TUCR transcription levels in comparison to transcription levels of known protein-coding and non-coding genes, in gliomas, as this has not been done. [56] To accomplish this, we first analyzed their spatial associations with markers for active chromatin (H3K4me3), active enhancers (H3K27ac), lncRNA transcription (RNA Pol.II) and open chromatin (ATAC-Seq). [56] We determined the significance of the spatial relationships between these marks and TUCRs utilizing publicly available U87 glioma cell CHIP- and ATAC-Seq datasets. Then, we compared the data to TUCR intervals that were randomly shuffled to create a negative control, other classes of non-coding RNAs, and TUCRs subset by genomic annotation (Supplementary Figure 2A). [56] We found that TUCRs displayed a significant enrichment for all transcriptional activity markers over expected and compared to control. Expected values are the chi-squared test’s predicted number of overlaps. The above data show that TUCRs are distributed throughout the genome, resistant to variation, and actively transcribed in U87 GBM cells. [56]

### TUCRs are highly expressed in GBM and LGG tumors

TUCR expression has not been characterized in GBM or LGG before. We performed the first comprehensive analysis of TUCR expression in these cancers by comparing GBM (n = 166) and LGG (n = 504) tumor samples from the Cancer Genome Atlas (TCGA) [33] to their normal brain cortex counterparts in TCGA (n = 5) and the Genotype-Tissue Expression Database (GTEx, n = 255). [34] We first analyzed absolute TUCR expression, as measured by transcripts-per-kilobase million (TPM). The absolute expression, in GBM, of all TUCRs was compared to the expression of lncRNAs, coding genes, antisense RNAs, and small noncoding RNAs (< 200 nt length), and the expression of TUCRs separated by genomic annotation into exonic, intronic, exonic/intronic, and intergenic (Supplementary Figure 3A). All gene annotations were obtained using the CHESS gene catalog, which contains most Refseq and Ensembl genes, while also including a series of understudied novel genes.[35] Highly expressed genes are visualized via heatmap (>=10 TPM, red) along with moderately (>=1 TPM, white) and lowly expressed genes (<1 TPM, blue). These analyses were repeated in LGG (Supplementary Figure 3B). The data show that intragenic TUCRs are highly expressed to a similar or greater degree as those of protein coding genes, in both GBM and LGG. These TUCRs may be an integral part of their “host genes,” while intergenic TUCRs demonstrate expression levels that are closer to those of lncRNAs and may instead represent novel transcripts of their own.

### TUCRs are deregulated in gliomas, and deregulation is associated with clinical outcomes

We analyzed TCGA tumor data and GTEx normal brain cortex data and found that in addition to being highly expressed in gliomas, TUCRs are highly deregulated in GBM and LGG as compared to normal brain cortex. As IDH-mutation status can be a confounding variable, we separated TCGA LGG samples into IDH wild-type (n = 382) and IDH1-mutant (n = 122) cohorts while removing all IDH1-mutant samples from the GBM (n = 8) cohort (Supplementary Figure 4). Of the 481 annotated TUCRs, we identified 95 that were upregulated and 69 that were downregulated in GBM (Figure 1A). We also identified 83 TUCRs that were upregulated and 50 TUCRs that were downregulated in IDH1-WT LGG (Figure 1B) and 53 upregulated and 70 downregulated in IDH1-mutant LGG (Figure 1C). The fact that there were many TUCRs that were differentially expressed in both IDH1-wildtype and -mutant LGG samples indicates that TUCRs are expressed and deregulated in an IDH1-independent manner. Of the 164 deregulated TUCRs in GBM, 59 were also deregulated in LGG, a 36% overlap (Figure 1D). Notably, there are multiple deregulated TUCRs across all TUCR annotation categories (Supplementary Figure 5A).

**Figure 1.**
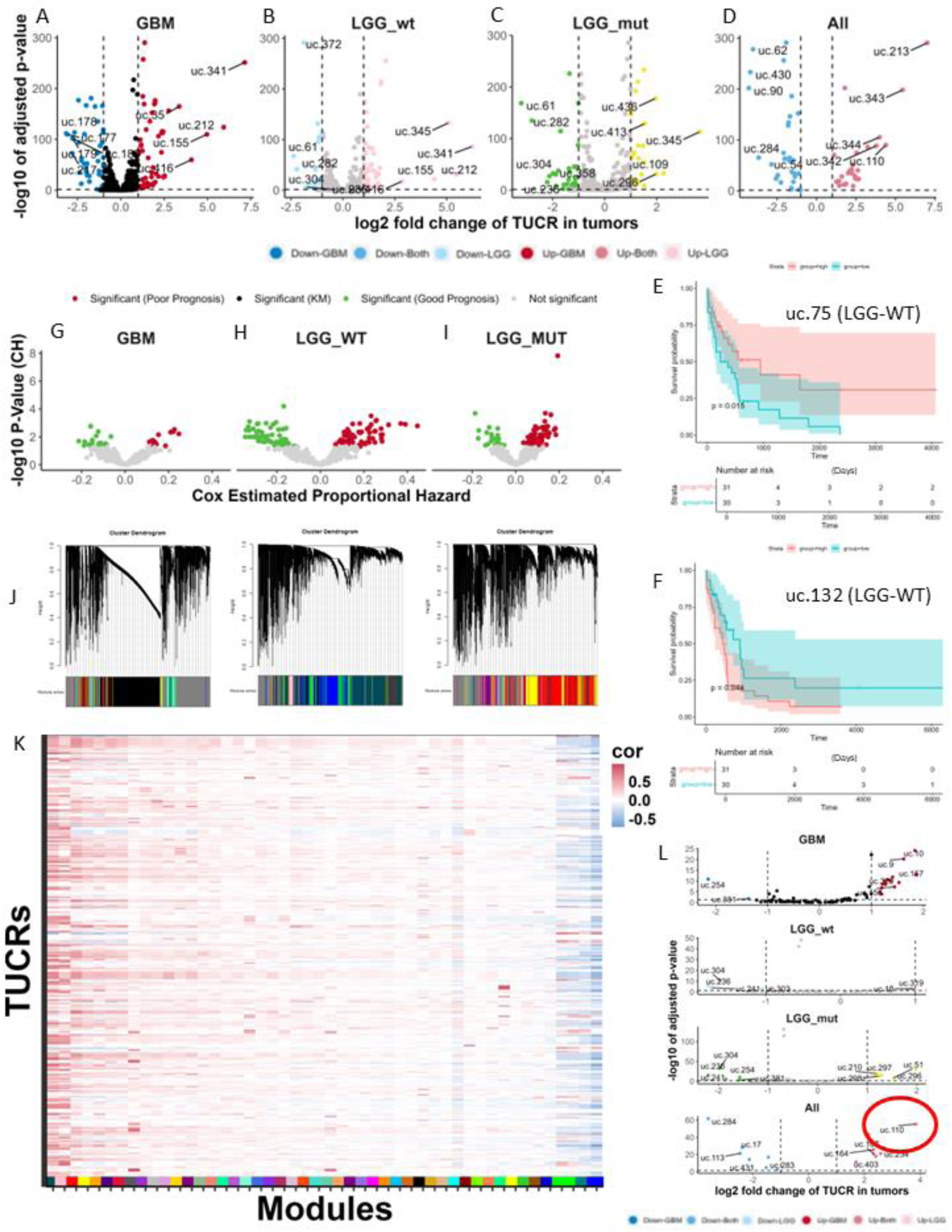
TUCRs are deregulated and associated with patient outcomes in GBM and LGG and may have broad functional roles. All experiments were performed using TCGA GBM and LGG RNA-Seq data. A) Volcano plot showing that 95 TUCRs are significantly upregulated (red) >=2-fold (1-log2FC) and 69 are significantly downregulated (blue) in GBM. B) Volcano plot showing that 83 TUCRs are significantly upregulated (pink) >=2-fold, and 50 are significantly downregulated (light blue) in IDH1 wild-type LGG. C) Volcano plot showing that 53 TUCRs are significantly upregulated (yellow) >=2-fold, and 70 are significantly downregulated (green) in IDH1 mutant LGG D) Volcano plot showing 29 upregulated (ash red) and 30 downregulated (medium blue) TUCRs across all conditions, which were included in the counts for A, B, and C. E) Kaplan-Meier showing that uc.75 is significantly associated with good prognosis in LGG-WT samples. Red line represents the uc.75 high expression group. Teal line represents the uc.75 low expression group. F) Kaplan-Meier showing that TUCR uc.132 is (green) patient prognosis across GBM, H) IDH1-wildtype LGG, I) and IDH1-mutant LGG. J) Gene similarity dendrograms from weighted gene correlation network analysis (WGCNA). 42,644 genes were aggregated into 3 “blocks” by gene similarity and were then further aggregated into 48 linkage modules using TUCR expression as trait data. Modules are denoted with distinct color hex codes. (e.g. #004C54 is the “midnight green” module) K) Heatmap showing that TUCRs demonstrate association with all 48 modules, suggesting broad potential functions. Red and Blue represent positive and negative correlations, respectively. L) Volcano plot showing that the uc.110 TUCR is the most upregulated TUCR in GBM and LGG and therefore may function as a novel oncogenic lncRNA. * = FDR < 0.05

We then sought to determine whether deregulation of TUCR expression correlates with patient outcomes in LGG-IDH-WT, LGG-IDH-MUT, and GBM (IDH-WT). For each of the 481 TUCRs, we generated a Kaplan-Meier (KM) plot tracking differences in survival for high expressing (upper quartile) and low expressing (lower quartile) tumor groups. We have highlighted two TUCRs that represent a statistically significant correlation with good (uc.75, Figure 1E) or poor (uc.132, Figure 1F) prognosis using both methods. The Bonferroni hypothesis adjusted p-value of the difference between groups was used for the determination of statistical significance. To further assess TUCR survival, we performed a separate workflow calculating the cox coefficient of the proportional hazard (CPH) of each TUCR, again assessing for statistical significance. Of the TUCRs that are expressed in GBM TCGA RNA-Seq data, relatively few were correlated with survival in a hypothesis adjusted, statistically significant, manner in our CPH workflow (FDR <= 0.05, Figure 1G). We considered that this low prevalence of survival associated TUCRs in GBM was due to the poor overall survival of GBM patients (∼15 months). We also studied survival differences in LGG patients, separated in IDH1-WT and IDH-MUT cohorts, as LGGs collectively have a longer median survival (∼84 months). Numerous TUCRs associated with survival both negatively and positively in LGG-WT and LGG-MUT (Figure 1H,I). While IDH1 mutation status can have an effect on patient prognosis, with IDH1-mutant patients generally having better overall survival, numerous TUCRs associate with survival both negatively and positively is not dependent on the IDH mutational status of the tumor.

When separated by annotation category, intragenic TUCR deregulation has a greater association with patient outcomes than intergenic TUCR deregulation (Supplementary Figure 5B). Detailed results for individual TUCRs can be found as supplementary materials (Supplementary Figures 6 and 7, TUCR Database).

### TUCRs are coregulated with genes that have known functions

We predicted TUCR functions by identifying coregulated genes with known functions via weighted gene co-expression network analysis (WGCNA).[36] We aggregated the 42,644 genes (Figure 1J) in our dataset into 46 colored modules based on clustered gene ontology (GO) terms. Each of these modules contains genes with known functions, such as RNA binding, cell signaling, immune response, metabolic response, etc. The data can also be used to predict gene function for novel genes, such as the TUCRs, by associating them with these genes that have known functions, as these associated genes are likely to share a function with the TUCR that they are associated with. To do this, we aggregated all 481 TUCRs into our modules. We identified TUCRs that correlate with each of the 46 modules, with some having positive correlations and others negative. (Supplementary Table 4) For example, TUCRs that exhibit a positive correlation with the #004C54 “midnight green” module (Supplementary Figure 8) could have a promoting effect on nucleic acid binding and regulation, while those that are negatively correlated with the #0000FF module (Supplementary Figure 9) may have a negative effect on ion channel signaling. Since many different TUCRs show associations with different modules, and every module has at least one TUCR that is associated with it, these results suggest that TUCRs have a broad range of potential functions in GBM and LGG (Figure 1K). WGCNA analyses were performed on all 481 TUCRs. Detailed results for individual TUCRs can be found as supplementary materials (Supplementary Figures 6 and 7, TUCR Database).

### TUCR, uc.110, is highly upregulated in gliomas and is predicted to bind nucleic acids

We utilized the expression and survival analyses to identify potentially important TUCRs in the context of gliomas. Of note, deregulated intergenic TUCRs may represent novel lncRNAs due to their similar expression levels, genomic location, and lack of coding potential.[2] These TUCRs are also easier to study experimentally; they are often thousands of kilobases (kb) from the nearest protein-coding gene and likely function in a manner that is independent of a “host gene”. Because of this, we focused on intergenic TUCRs for our experimental studies. Of the deregulated intergenic TUCRs in GBM and in LGG (Figure 1L). We found that uc.110 is the most upregulated as compared to normal brain; 21-fold in GBM and 53-fold in LGG (Figure 2A-2B). It has near binary expression; it is rarely expressed at all in normal brain but is very highly expressed in GBM and LGG (Figure 2C-2D). It does not appear to have a correlation with patient outcomes. (TUCR Database, uc.110). Despite this, due to its high deregulation, we hypothesized that this TUCR is functioning as a tumor promoter, as many known GBM driver genes also have weak or insignificant correlations with patient outcomes (Supplementary Table 5).

**Figure 2.**
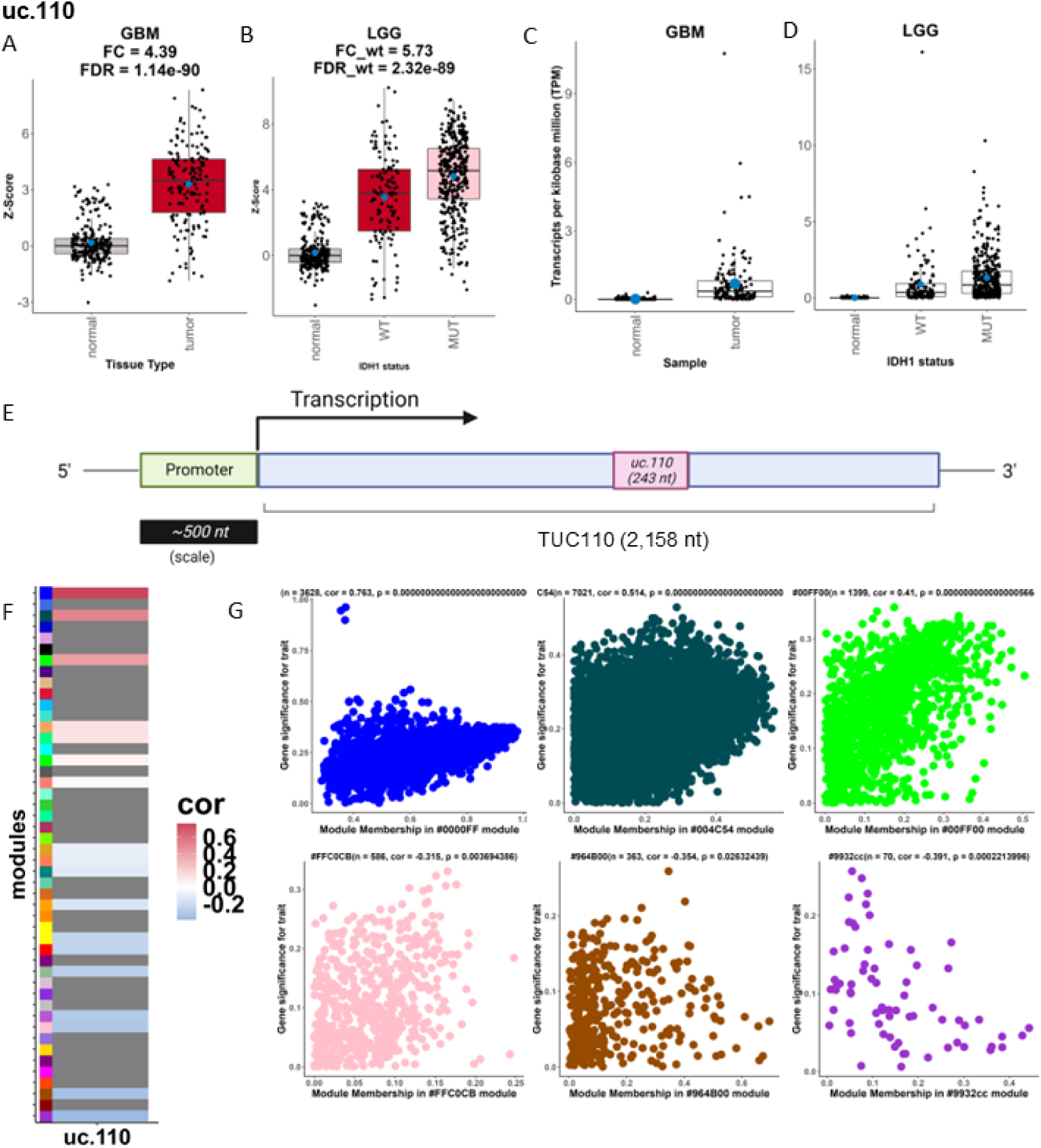
The uc.110 TUCR is the most upregulated intergenic TUCR in gliomas. A) Box- and dotplot showing that uc.110 is 21-fold upregulated in GBM and B) 53-fold upregulated in LGG based on TCGA data analysis. Red boxes represent upregulated TUCRs. Gray boxes represent TUCRs that are not deregulated. C) Box- and dotplot showing that uc.110 is expressed in tumors but is poorly expressed in normal brain cortex based on GBM and D) LGG TCGA and Gtex data analyses. E) Cartoon showing that uc.110 is a 243 nt region in a 2,158 nt transcript. F) Heatmap depicting uc.110 gene module association. Positive correlations are red, while negative correlations are blue, with weak correlations in white. Modules with no linkage are gray. G) Scatter plots depicting uc.110 association with top 3 positive (top row) and negative (bottom row) correlation modules. Caption lists the module name, number of genes in the module, and the significance of uc.110 association with the module. (Similar figures for other putative oncogenic TUCRs available at in the TUCR Database provided as supplementary materials)

Since many TUCRs exist as a part of a larger transcript [2], we first determined the sequence of the uc.110 full transcript. We utilized machine learning and *de novo* transcript reassembly using TCGA and GTEx RNA-seq data to reconstruct RNA-Seq transcripts in the absence of a reference genome (Supplementary Figure 10A). [35] We identified a 2,158 nucleotide (nt) long RNA molecule that contains the 243 nucleotide (nt) uc.110 ultraconserved sequence (Figure 2E) as a novel transcript. We confirmed the existence of this transcript experimentally using PCR amplifications and sequencing (Supplementary Figure 10B).

After identifying the sequence for the full uc.110 transcript (Supplementary Figure 10C), we used qPCR to confirm its independence from its closest protein coding gene, GBX2. We found that knock-down of uc.110 with two different siRNAs did not alter the expression of GBX2 in GBM cell lines, thus demonstrating that uc.110 and GBX2 are two separate transcripts (Supplementary Figure 10D-10E). Based on the work by Mestdagh et al [56], we utilized our WGCNA workflow to identify genes and modules (Figure 2F) that are significant to this transcript. Of note, one of the top modules for uc.110 by module association is the #004C54 module, which represents genes that are involved in nucleic acid and protein binding (Supplementary Figure 8). Genes that are members of these modules are positively coregulated with uc.110 (Figure 2G). Based on these findings, we hypothesized that uc.110 may be operating as a tumor promoting RNA-binding molecule. We also performed similar analyses for all 481 TUCRs to identify potential functional roles for each TUCR in gliomas. Examples of a tumor promoting TUCR (uc.2, Supplementary Figure 6, TUCR Database) and a tumor suppressive TUCR (uc.15, Supplementary Figure 7, TUCR Database) are depicted in this manuscript, while the rest of the 481 TUCRs are shown in the supplementary materials (TUCR Database).

### uc.110 has tumor promoting effects in GBM

To determine the function of uc.110 in GBM, we first used qPCR to investigate the expression of uc.110 using RNA in our banked tumor samples compared to normal brain cortex and cell lines compared to normal human astrocytes. We independently confirmed the results from our TCGA analysis showing uc.110 is highly upregulated in GBM tumors (Figure 3A, 3B). We then designed two siRNAs that target separate regions on the uc.110 RNA, one that begins at nucleotide 96/243 (si-uc.110-1) and one that begins at nucleotide 195/243 (si-uc.110-2), as well as a scrambled control (si-SCR) (Supplementary Figure 11A). We generated stable A172 and U251 GBM cell lines that express uc.110 (LV-uc.110) or the empty expression vector (LV-pCDH). We subjected these cell lines to siRNA transfection and assessed the effects on cell counting, survival and invasion assays (Supplementary Figure 11B). We used qPCR to show that uc.110 is generally, though not uniformly, upregulated in GBM cells (Supplementary Figure 11C,10D,10E). Based on these data, we prioritized the use of cell lines that overexpress uc.110 (A172, U251) for knockdown experiments, and cells that express lower levels of uc.110 (U87, GSC-28) for overexpression experiments. We confirmed that siRNA targeting of uc.110 lead to knockdown of uc.110 expression in A172 and U251 cells (Figure 3C). We also confirmed that LV-uc.110 overexpresses uc.110, and that this overexpression rescues uc.110 bioavailability in A172 and U251 cells (Figure 3C).

**Figure 3.**
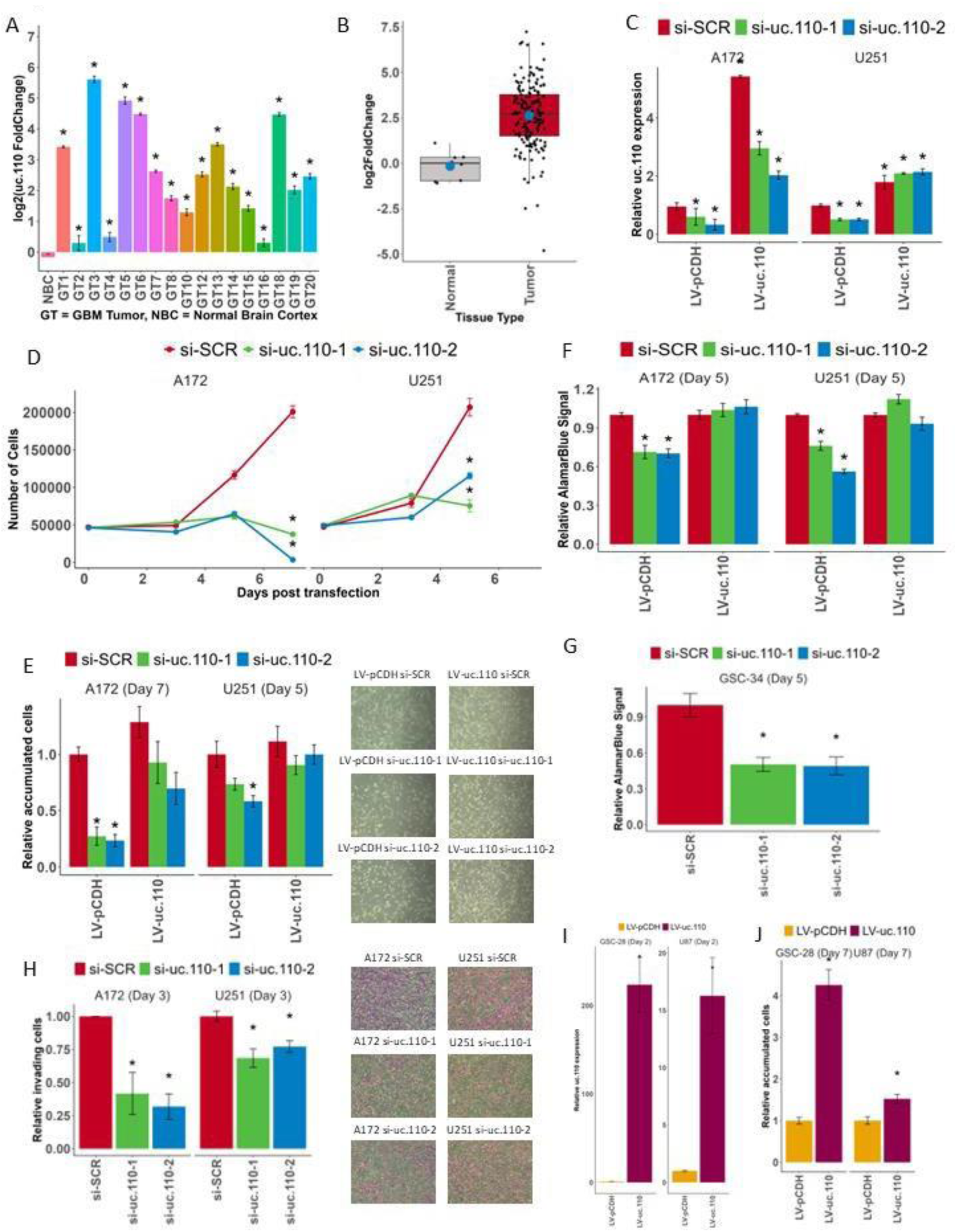
Uc.110 TUCR operates as a tumor promoter gene. A) Bar graph depicting uc.110 upregulation in banked UVA GBM tumors (GT) versus normal brain cortex (NBC). B) Boxplot representing uc.110 expression in pooled tumors versus normal brain. Red boxes indicate an upregulated TUCR. C) Bar graph depicting qPCR validation of uc.110 siRNA knockdown and rescue in A172 and U251 cell lines. si-SCR = scrambled control siRNA (red), si-uc.110-1 = siRNA targeting uc.110 at nucleotide 96/243 (green), si-uc.110-2 = siRNA targeting uc.110 at nucleotide 195/243 (blue). D) Line graph showing that knockdown of uc.110 reduces A172 and U251 cell accumulation over a 5-7 day period. E) Bar graph depicting that the cell accumulation phenotype is rescued when uc.110 is overexpressed in the presence of siRNA. Images are representative of the listed sample. F) Bar graph showing that knockdown of uc.110 reduces A172 and U251 cell viability via Alamar Blue assay and can be rescued with uc.110 overexpression. G) Bar graph showing that knockdown of uc.110 reduces GSC-34 glioma stem cell viability via AlamarBlue. H) Bar graph showing knockdown of uc.110 reduces A172 and U251 cell invasion and migration. Images are representative of the listed sample. I) Bar graph showing the qPCR validation of overexpression of uc.110 in GSC-28 glioma stem cells and U87 cells. J) Bar graph showing that overexpression of uc.110 increases cell accumulation in U87 and GSC-28 cells. Facets represent cell lines. * = p < 0.05

Next, we performed cell counting assays [20, 37–39] to determine the effects of uc.110 knockdown and rescue on cell accumulation. When we reduced uc.110 expression, we reduced cell accumulation in A172 and U251 cells (Figure 3D). When we rescue uc.110 bioavailability, the cell accumulation phenotype is restored in A172 and U251 cells (Figure 3E). We then used Alamar Blue [40, 41] to measure cell viability. When we reduce uc.110 expression, A172 and U251 cell viability is reduced. We were able to rescue this phenotype by increasing uc.110 bioavailability (Figure 3F). We observed a similar phenotype in a glioma stem cell line that overexpresses uc.110 (GSC-34, Figure 3G).

We then investigated the invasive potential of uc.110 using a transwell invasion assay. [42–44] Knockdown of uc.110 reduced A172 and U251 cell invasion through a collagen IV matrix (Figure 3H). When uc.110 bioavailability was increased, a partial recovery of the phenotype is observed (Supplementary Figure 11F). Lastly, we overexpressed uc.110 in U87 and GSC-28 cells (Figure 3i) and determined that this leads to increased cell accumulation compared to the empty vector after 7 days (Figure 3J) in U87 and GSC-28 cells. These data show that uc.110 has tumor enhancing effects in GBM cells and stem cells.

After determining that uc.110 displays a tumor promoting phenotype *in vitro*, we sought to determine whether this effect is recapitulated *in vivo*. U251 GBM cells were transfected with si-uc.110-1 or si-uc.110-2. After 2 days, these cells were implanted into immunodeficient mice using intracranial injection (Supplementary Figure 12A). [37, 38, 45, 46] Tumor growth was monitored by MRI and mouse survival was observed over a period of 70 days. Mice that were xenografted with U251 cells that were transfected with si-uc.110-1 and si-uc.110-2 expression developed smaller tumors, as depicted, and quantified by MRI (Figure 4A, 4B). The mice that received si-uc.110 also displayed better overall survival than mice that received scrambled control siRNA cells (Figure 4C).

**Figure 4.**
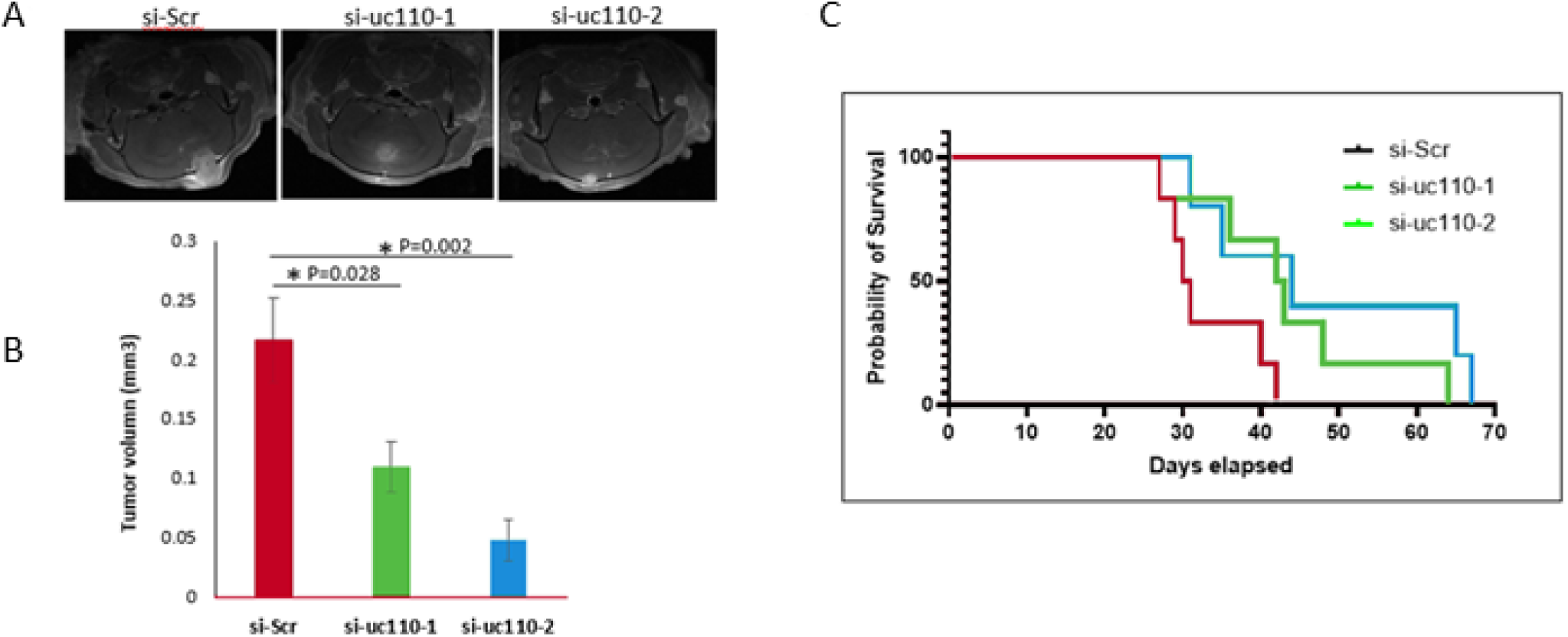
The uc.110 TUCR promotes tumor growth in vivo. A) MRI images reveal a reduction in tumor size when uc.110 is knocked down via siRNAs. B) Bar graph showing that knockdown of uc.110 leads to a reduction in tumor volume. si-SCR = scrambled control siRNA (red), si-uc.110-1 = siRNA targeting uc.110 at nucleotide 96/243 (green), si-uc.110-2 = siRNA targeting uc.110 at nucleotide 195/243 (blue). C) Kaplan-Meier plot showing that knockdown of uc.110 leads to increased mouse survival.

### uc.110 regulates the expression of the Wnt pathway member, Membrane Frizzled Related Protein (MFRP)

LncRNAs can have various functions that depend on their subcellular localization. Nuclear lncRNAs are usually involved in transcriptional regulation, while cytosolic lncRNAs are usually involved in translational and spatial regulation. [2] We fractioned four GBM cell lines (A172, U251, U87, U1242) into nuclear and cytosolic fractions. When compared to nuclear (U44, U48) and cytosolic (GADPH, PPIA) controls, uc.110 appears to be localized to both the nucleus (mainly in U87, U251, and U1242 cells), and the cytoplasm (mainly in A172 cells) (Supplementary Figure 11G). We then performed RNA-Seq on A172 cells that had been transfected with si-SCR, si-uc.110-1, or si-uc.110-2 for 48 hrs. and found several genes that are deregulated when uc.110 expression is downregulated (Figure 5A). To identify genes that are particularly related to uc.110 function, we focused on genes that demonstrated coregulation with uc.110 in our WGCNA analysis (Figure 2F). Of particular interest was the membrane frizzled related protein, also known as MFRP. [47, 48] MFRP serves as a shuttle for the Wnt-ligand, and functions as an activator of the Wnt-signaling pathway. This gene was the only gene in our analysis that correlated with uc.110 expression, was upregulated in GBM tumors, and downregulated when uc.110 is knocked down in A172 cells, suggesting MFRP coregulation with uc.110. (Figure 5B). Notably, there is currently no literature on uc.110 in gliomas, no published literature on its relationship with MFRP, and no published literature on TUCR associations with MFRP.

**Figure 5.**
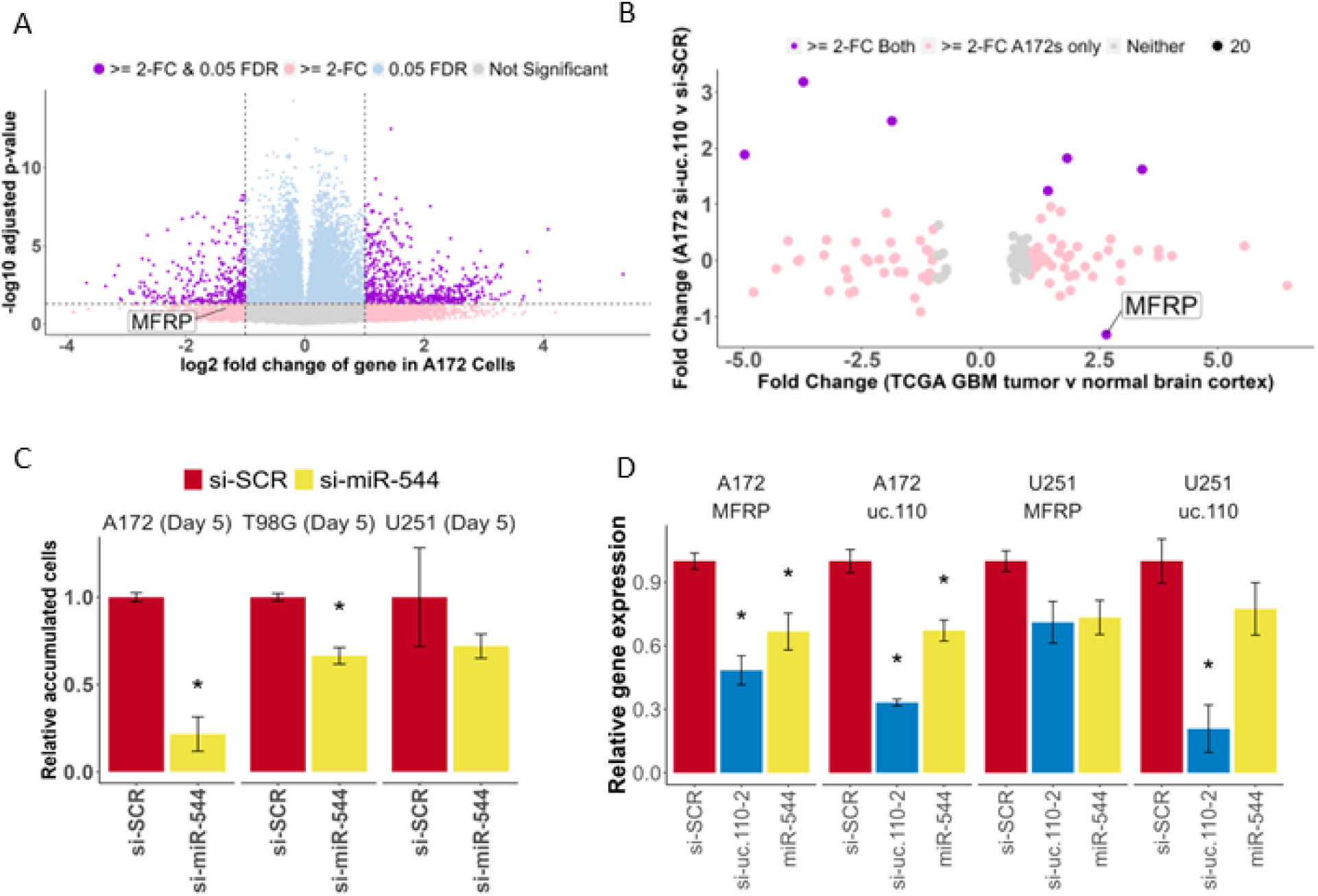
The uc.110 TUCR activates Wnt-signaling by sponging miR-544 from membrane frizzled related protein (MFRP) 3’UTR. A) Volcano plot depicting transcriptome deregulation in RNA-Seq data on A172 cells transfected with si-uc.110. Purple dots represent genes that are significantly deregulated >= 2-fold. Blue dots represent genes that are significantly deregulated, but with a FC <2. Pink dots represent genes that are deregulated >= 2-fold, but are not statistically significant (FDR). Gray dots are neither deregulated nor significant. B) Dot plot showing that, of the genes that predicted miR-544 targets, MFRP is the only gene that is upregulated in GBM Tumors and downregulated when uc.110 is downregulated. Purple dots represent predicted miR-544 targets that are deregulated in A172 cells from 5A and TCGA RNA-Seq data. Pink dots represent predicted miR-544 targets that are deregulated in A172s from 5A only. C) Bar graph showing that miR-544 transfection reduces cell accumulation in A172, T98G, and U251 cells, confirming its tumor suppressor role. Facets represent cell lines. si-SCR = scrambled control siRNA (red), si-uc.110-2 = siRNA targeting uc.110 at nucleotide 195/243 (blue). miR-544 = miR-544 (yellow). D) Bar graph showing that transfection with miR-544 or si-uc.110-2 reduces uc.110 and MFRP expression.

### uc.110 sponges the tumor suppressor microRNA miR-544 to increase the bioavailability of MFRP and WNT activity in GBM

One common lncRNA mechanism of action is as a miRNA sponge, acting as a binding competitor for various miRNAs and therefore increasing the bioavailability of those miRNAs’ targets. [2, 49, 52–53] Based on the WGCNA data that we generated above, we hypothesized that uc.110 may function by sponging miRNAs away from MFRP transcripts, as their expression relationship is consistent with such an interaction. We hypothesized that a tumor suppressor miRNA can successfully target and suppress MFRP in the normal brain (Supplementary Figure 14A). This leads to downstream activation of Wnt target genes involved in biological processes such as cell accumulation, invasion, and stem cell differentiation (Supplementary Figure 14B).

We further hypothesized that in glioma tumors, uc.110 is activated and acts as a binding competitor for this miRNA (Supplementary Figure 14C), increasing the bioavailability of MFRP and increasing Wnt pathway signaling (Supplementary Figure 14D). To identify candidate miRNAs that are consistent with the afore mentioned hypothesis, we screened public databases and published literature for GBM tumor suppressor miRNAs that are predicted to bind to both uc.110 and MFRP. The only miRNA that fulfilled these criteria was miR-544. We first investigated the functional effects of miR-544 in GBM cells. Transfection of miR-544 into U251, A172, and T98G GBM cell lines reduced cell accumulation after 5 days (Figure 5C). The use of an additional GBM cell line, T98G, for these experiments was to further validate its previously published role as a tumor suppressing miRNA. Expression of both uc.110 and MFRP was statistically significantly reduced when A172 cells transfected with miR-544 or si-uc.110 (Figure 5D). U251 cells demonstrated a similar reduction in expression for both genes, but the reduction was not statistically significant.

To further test the hypotheses, we asked if miR-544 targets both uc.110 and MFRP, and if this binding affects Wnt signaling. To determine whether MFRP and uc.110 are direct targets of miR-544, we constructed luciferase reporter vectors by inserting the uc.110 ultraconserved region and MFRP 3’UTR downstream of hRluc followed by Synthetic Poly(A) using psiCHECK-2 backbone vector (Promega) (Figure 6A,6B). We first measured target binding by transfecting the reporter constructs followed by transfection with miR-544 or miR-SCR (control) in GBM cells. Ectopic expression of miR-544 significantly decreased luciferase activity compared to miR-SCR (Figure 6D, left panel and figure 6E, left panel). These binding sites for miR-544 were predicted via computational algorithms and validated via sequencing. We then mutated the binding sites for MFRP and uc.110 (Supplementary Figure 13, Figure 6C) and assessed signal strength again. The data showed that luciferase activity was not significantly altered in mutant-reporter-vectors transfected cells (Figure 6D, right panel and Figure 6E, right panel), indicating that miR-544 binds to both uc.110 and MFRP in GBM cells, and that this binding is lost when the miRNA binding sites are mutated.

**Figure 6.**
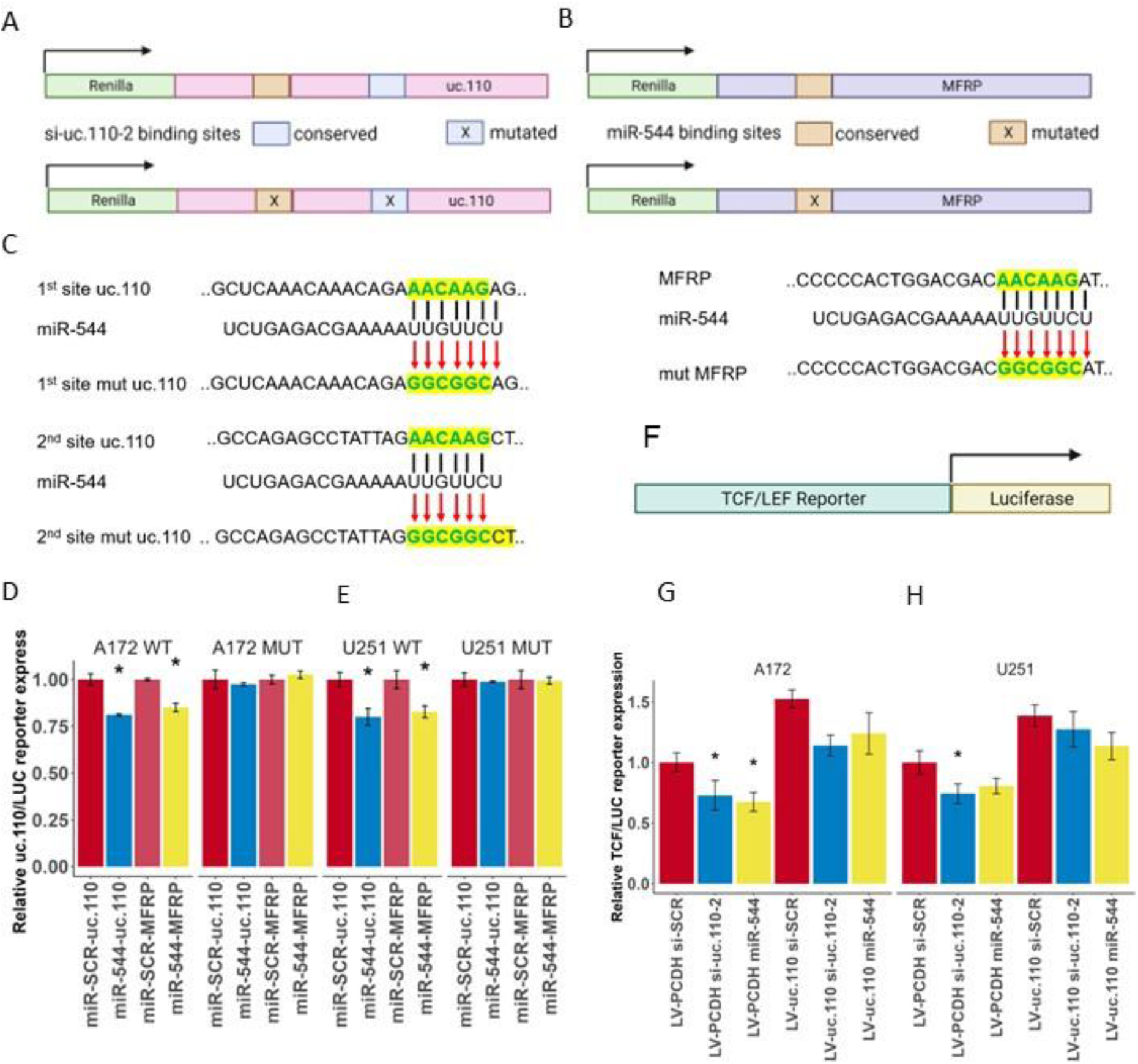
The uc.110 TUCR activates Wnt-signaling by sponging miR-544 from membrane frizzled related protein (MFRP) 3’UTR. A) Schematic depicting the uc.110 luciferase construct used to demonstrate binding to miR-544 and si-uc.110-2. Binding of miR-544 to binding sites (orange) leads to a degradation of construct and a reduction in Renilla signal (Green) B) Schematic depicting the MFRP luciferase construct used to demonstrate binding miR-544. C) Schematic depicting miR-544 binding site mutations for uc.110 (two sites) and MFRP (one site). Top row represents wild-type binding sites. Middle row is the miR-544 binding region. Bottom row are mutated sites. D) Bar graph showing that transfection of miR-544 reduces uc.110 and MFRP luciferase expression signal in A172 and E) U251 glioma cells, and that mutating miR-544 binding sites rescues luciferase signal. Facets represents cell lines and miR-544 binding site mutation status. F) Schematic depiction of TCF/LEF luciferase reporter construct used to measure downstream Wnt-signaling pathway activation. TCF/LEF binds to the reporter region (green) and activates luciferase (yellow). G) Bar graph showing that transfection of si-uc.110-2 and miR-544 reduces TCF/LEF reporter signal in A172 and H) U251 glioma cells. Signal is rescued when uc.110 is overexpressed in the presence of siRNA or miR-544. * = p< 0.05

Lastly, we asked if uc.110 expression alters Wnt pathway activity. To answer this question, we studied one of the most established downstream targets of Wnt-signaling, the T cell factor/lymphoid enhancer factor family (TCF/LEF). When Wnt-signaling is activated, TCF/LEF is produced downstream and activates Wnt-signaling target genes. Therefore, TCF/LEF activity can be used as a proxy for pathway activity and can be measured with a TCF/LEF luciferase reporter assay. [57–58] (Figure 6F) The activity of this reporter can be regulated by either directly reducing Wnt bioavailability with miR-544 or indirectly by targeting uc.110 with siRNA. If upstream Wnt signaling is reduced, the luciferase construct will bind fewer activators and exhibit decreased signal. Likewise, we would expect that overexpression of uc.110 would rescue the bioavailability of MFRP and consequently also downstream activation of the TCF/LEF construct. We found that transfection of A172 and U251 cells with si-uc.110 and miR-544 reduced reporter activity in A172 (Figure 6G) and U251 (Figure 6H) cells, and that this effect can be rescued via uc.110 overexpression. These data taken in conjunction provide strong support for a miRNA sponge model for the uc.110 tumor enhancer. Altogether, the above data demonstrate an important role for uc.110 in regulating the Wnt pathway in GBM by sponging the Wnt inhibitory miRNA miR-544 (model shown in Supplementary Figure 14).

## DISCUSSION

This study investigated Transcribed Ultraconserved Regions (TUCRs), a set of transcripts that might contain long noncoding RNA sequences that are fully conserved across human, mouse, and rat genomes. These TUCRs are distinct due to their exceptional conservation, which often signifies functional importance. Despite their potential significance, TUCRs have been minimally explored, especially in relation to cancer. In fact, a PubMed search of TUCR, UCR, or “ultraconserved” and Cancer reveals only ∼ 70 publications in cancers but none in gliomas. Other classes of RNA, such as MicroRNAs (miRNAs) contain over 12,000 publications in cancer. Even single protein coding genes, such as TP53/p53, contain over 24,000 publications in cancer. [2] Of note, the findings of this study represent the first of their kind on TUCRs *in gliomas*. They contribute critical new insights into an uncharted area of glioma biology, while also providing a novel framework for studying TUCRs in other cancers and other diseases, where they are also understudied.

We confirmed that TUCRs are located across the genome, and showed that they are resistant to variation, and actively transcribed in U87 GBM Cells. We manually annotated each as either exonic, intronic, exonic/intronic, or intergenic. We identified distinct signatures for intergenic and intragenic (exonic, intronic, exonic/intronic) RNAs. Intragenic TUCRs are expressed at a level that is most like coding genes, and may be coded as part of their host gene transcripts, as they are less likely to be host gene independent. [56] Intergenic TUCRs, on the other hand, more closely resemble lncRNAs in terms of expression and may represent novel transcripts of their own, as they exist outside of their nearest gene.

We then performed the first analysis of TUCR expression in gliomas and found that the majority of TUCRs are deregulated >= 2-fold in GBM and LGG, with a 36% overlap. This shows that TUCRs are not only expressed, but also frequently dysregulated in gliomas compared to normal brain tissue. This is critical, as their high degree of conservation and dysregulation suggests that they may serve critical biological functions. For example, TUCRs that are associated with the critical exon 2 of various Hox genes, such as uc.212 (HOXA2) and uc.151 (HOXC10) are highly upregulated in gliomas, but it is unclear whether these genes are independent of their host transcripts. We then extended our analysis to TUCR correlation with patient survival. In GBM, the extremely short survival times (15 months) limit the detection of significant correlations. However, patients with LGG live substantially longer (84 months), and therefore more TUCRs are associated with patient outcomes in this disease, suggesting a potential impact on glioma patients’ prognoses and indicating possible novel biomarkers. Additionally, we found many TUCRs that were deregulated across annotation status and IDH1-status, which suggests that, annotation status may have an effect on absolute TUCR expression, neither of these additional factors had a substantial effect on TUCR deregulation or association with patient outcomes.

Another facet of our research involved predicting the functions and mechanisms of action of TUCRs in gliomas. We studied this for the first time in gliomas WGCNA workflows to cluster TUCRs and provide functional predictions based on shared functions between coregulated genes. This approach identifies a wide range of potential functions for TUCRs, encompassing activities such as nucleic acid binding regulation, stem cell differentiation, organ development, immune response, and cell signaling. This analysis focused on the associations of individual TUCRs with the functional modules, but it would also be worth studying these TUCRs as clusters of coregulated genes, perhaps through a high throughput single-cell approach.

We found intergenic TUCRs to be of notable interest because they resemble lncRNAs but are much more highly conserved and experience less sequence variation. Notably, these TUCRs do not overlap with known genes, suggesting they might represent novel lncRNAs. Of these TUCRs, uc.110 is the most upregulated in both GBM and LGG. Knocking down uc.110 reduces cancer cell characteristics *in vitro* and *in vivo* and improves survival in mouse models. On the other hand, increasing uc.110 expression increases malignancy in cells that do not express it, further indicating its potential tumor promoting role. We explored uc.110’s function via WGCNA, revealing its membership in modules associated with tumor enhancing nucleic acid binding. We integrated these data with transcriptome deregulation data (RNA-Seq) post-uc.110 knockdown, revealing a close relationship between uc.110 and the oncogenic membrane frizzled-related protein (MFRP). This protein is involved in activating the Wnt-signaling pathway, impacting cell proliferation, invasion, migration, and stem cell differentiation. Aberrant expression of Wnt-signaling is demonstrated in gliomas. [59–60] Therefore, we hypothesized that uc.110 might sponge tumor suppressor miRNAs from MFRP, enhancing Wnt signaling activation. Accordingly, we demonstrated that one mechanism of action for the uc.110 tumor promoter is as a miRNA sponge for miR-544, therefore increasing the bioavailability of MFRP. This is a novel interaction between all three genes as the published literature on the role of miR-544 and its effects on MFRP alone is non-existent, and likewise there is no published literature considering uc.110’s role in potentially mediating this interaction as a sponge. We also note that this is a likely cytosolic function for uc.110, as although miRNAs may target nuclear genes during cell division, the sponging model is generally considered to be a cytosolic event. Per the same WGCNA analysis, it is entirely possible that uc.110 has a separate nuclear function, perhaps in transcription factor binding (Figure 3, Supplementary Figure 8). This is a known function for lncRNAs, and uc.110 could therefore be operating in a similar way. We anticipate exploring this in future work, especially as uc.110 is shown to localize to the nucleus in some GBM cell lines. (Supplementary Figure 11G)

In conclusion, our results suggest that TUCRs are an important class of regulatory RNAs. They are more highly conserved than typical genes and more resistant to variation, which suggests biological importance. They are perturbed in gliomas, and this perturbation is associated with clinical outcomes. Our predicted functions reveal that TUCRs are widely involved in cancer-related biological processes. Some TUCRs previously thought to be intergenic may represent previously undiscovered genes. Our findings also identify and characterize uc.110 as a new tumor enhancer and likely oncogene in gliomas. Each of the experiments performed in our study represents the first of its kind in gliomas. We have developed, adapted, and presented novel methods for studying TUCRs that can be used in other cancers and other diseases, where TUCRs remain very understudied. These methods and the data derived from them represent a “TUCR database” that will serve the scientific community in future TUCR studies in gliomas and other diseases, where they remain unstudied or understudied.

## SUPPLEMENTARY DATA

**Supplementary Figure 1.**
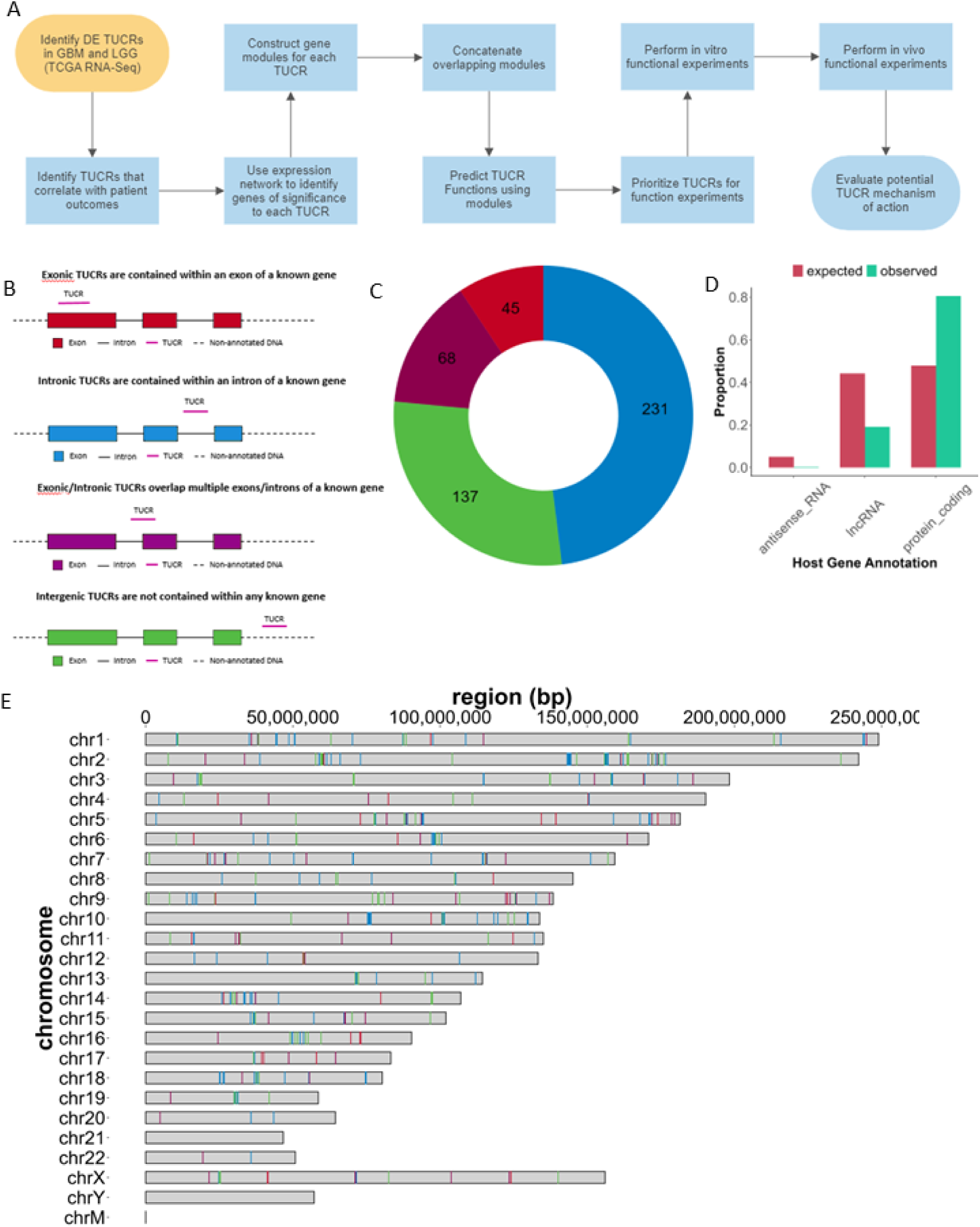
Annotation, localization and expression of TUCRs in GBM and LGG. A) Experimental workflow for identifying and studying TUCRs of interest. B) Genomic annotation shows that TUCRs can be exonic (red), intronic (blue), exonic/intronic (purple) or intergenic (green). C) Circle graph showing the distribution of genomic annotations across all 481 TUCRs, with colors matching. D) Bar plot depicting the classes of “host genes” that TUCRs associate with, showing an enrichment for protein coding genes. 1B. E) Karyoplot showing that TUCRs exist on all chromosomes except for the Y chromosome and mitochondrial DNA, vertical lines show TUCRs with colors matching 1B.

**Supplementary Figure 2.**
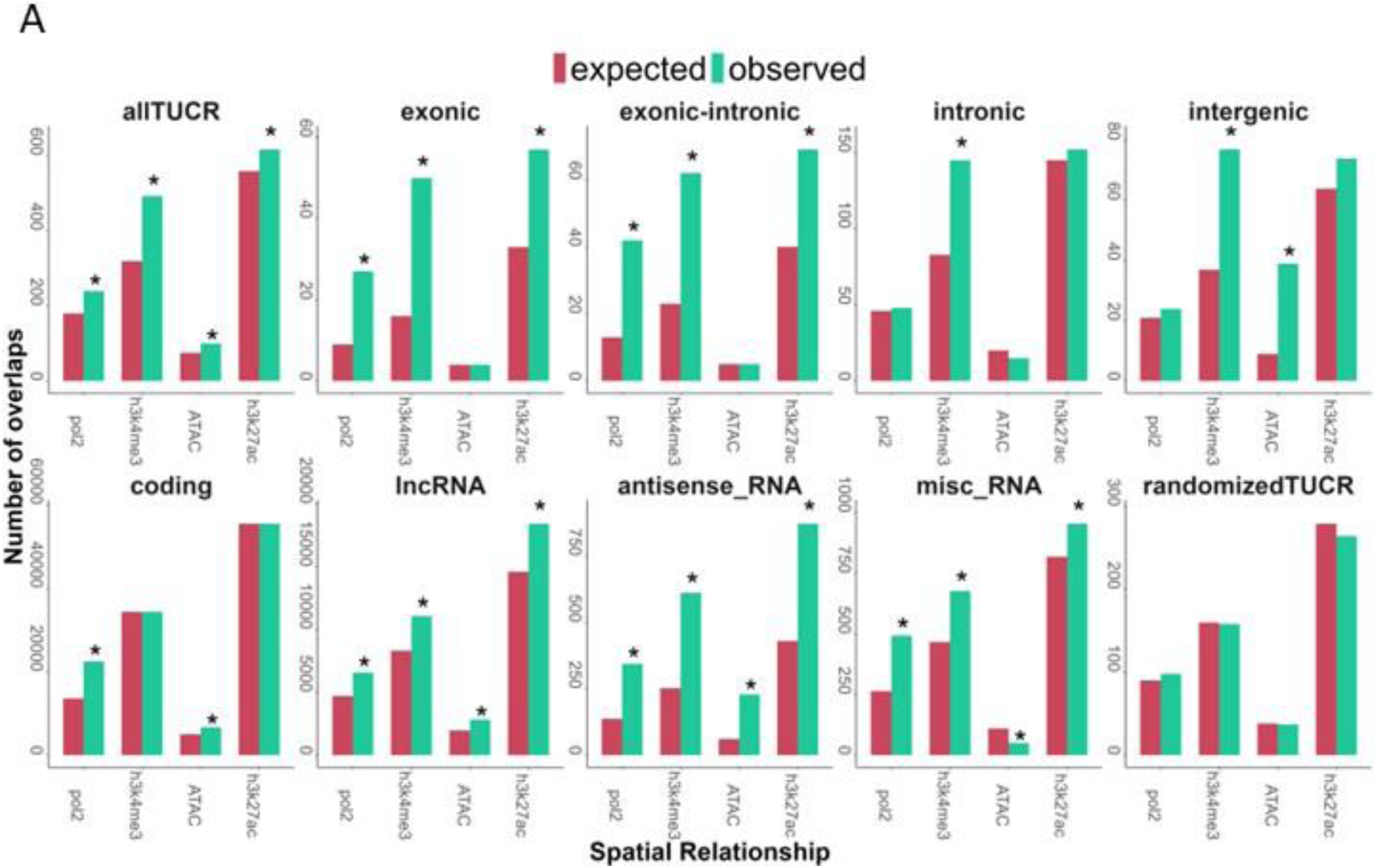
Annotation, localization and expression of TUCRs in GBM and LGG (cont.). A) Bar chart showing that TUCRs are enriched for markers for open and active chromatin in GBM U87 cells, suggesting that they represent transcriptionally active sites. Red bars represent chi-square expected overlaps, and teal bars represent observed values. * = FDR < 0.05

**Supplementary Figure 3.**
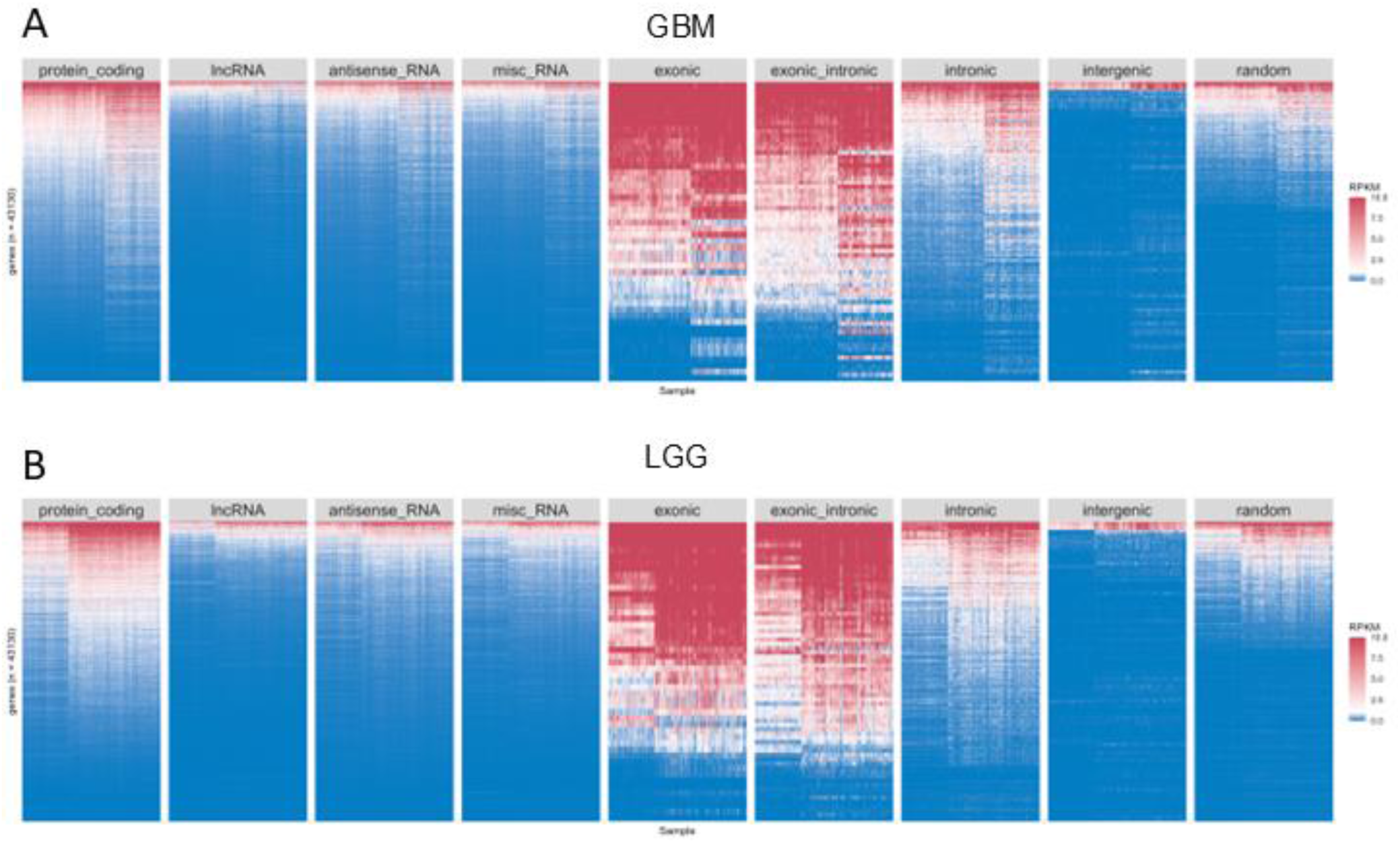
Annotation, localization, and expression of TUCRs in GBM and LGG. A) Heatmap representing TUCR absolute expression (TPM) across multiple gene annotations in GBM and B) LGG. Blue represents poorly expressed genes (<1 RPKM), White/Pink genes are moderately expressed (>=1 TPM) and Red represents highly expressed genes (TPM >=10). TUCRs demonstrate an expression profile that is comparable with protein coding genes. * = p < 0.05

**Supplementary Figure 4.**
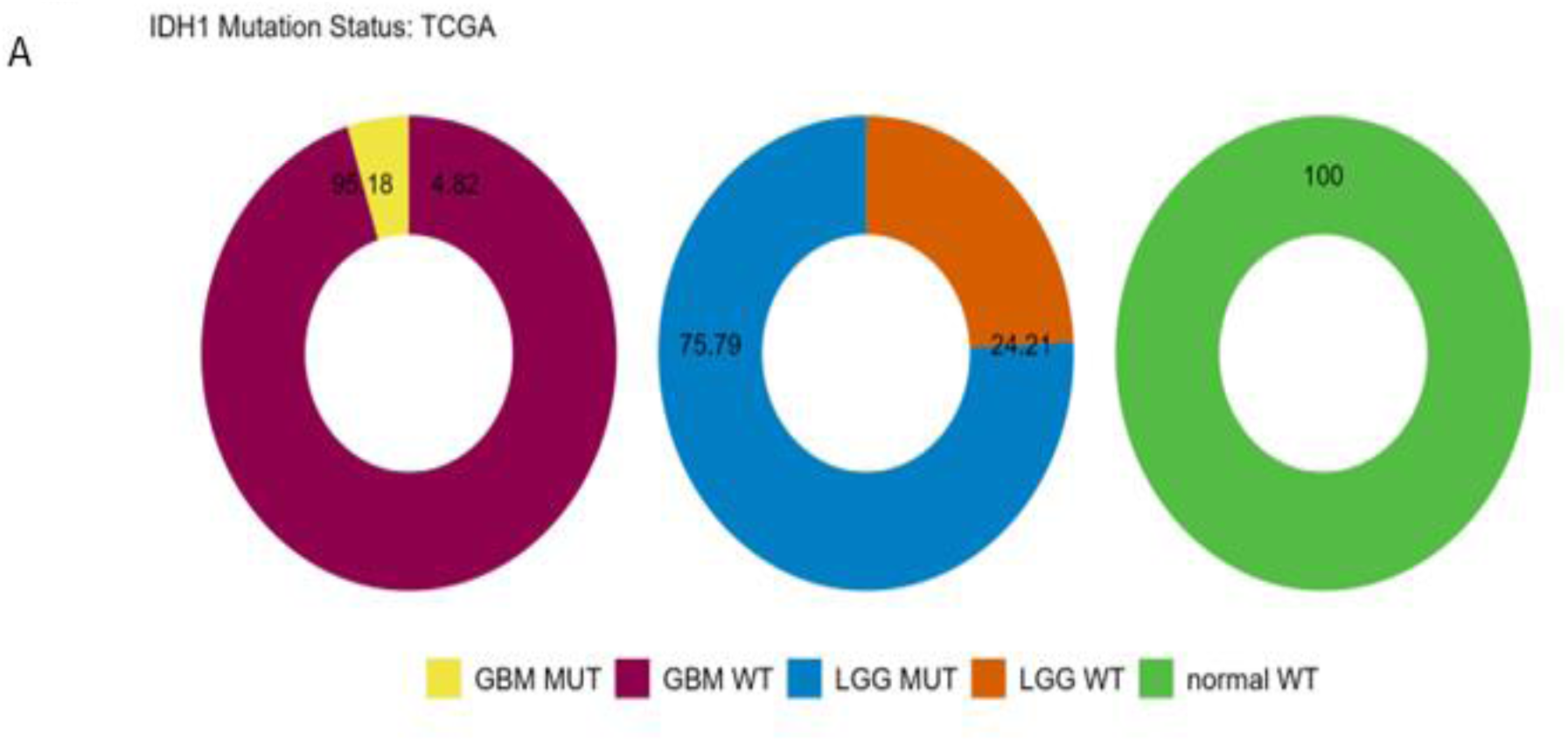
TUCRs are deregulated and correlated with patient outcomes in IDH1 mutant TCGA glioma samples A) Ring chart showing distribution of glioma samples by IDH1 mutation status. All experiments were performed using TCGA GBM and LGG RNA-Seq data.

**Supplementary Figure 5.**
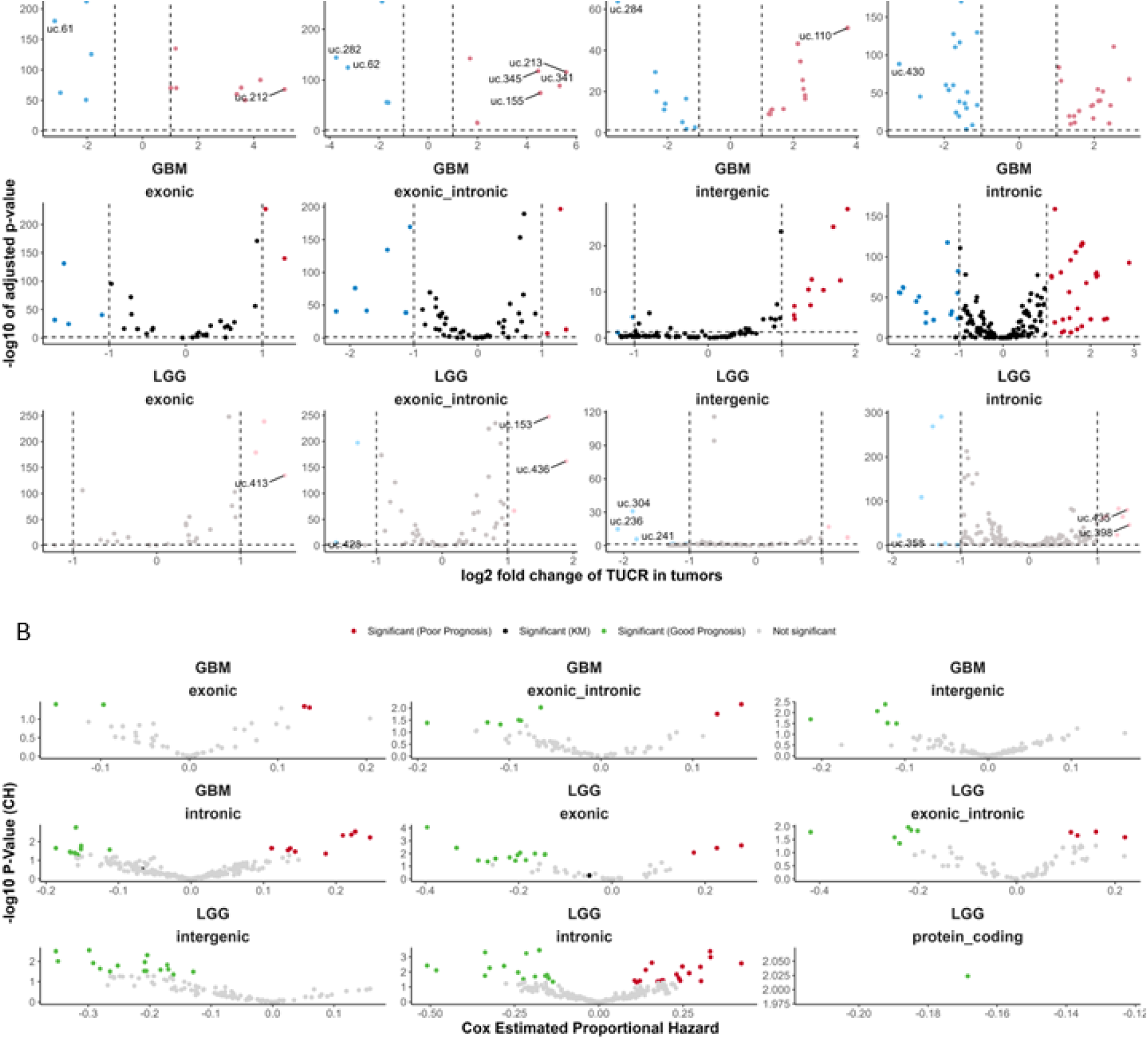
TUCRs are deregulated and associated with patient outcomes in gliomas and may have broad functional roles (cont.) All experiments were performed using TCGA GBM and LGG RNA-Seq data. A) Volcano plots showing that TUCRs are deregulated in every TUCR annotation category in GBM and LGG. Red dots are upregulated. Blue are downregulated. B) Volcano plot showing that TUCRs in every TUCR annotation category are associated with patient outcomes in gliomas. Red dots represent TUCRs significantly associated with poor prognosis. Green dots represent TUCRs significantly associated with good prognosis.

**Supplementary Figure 6.**
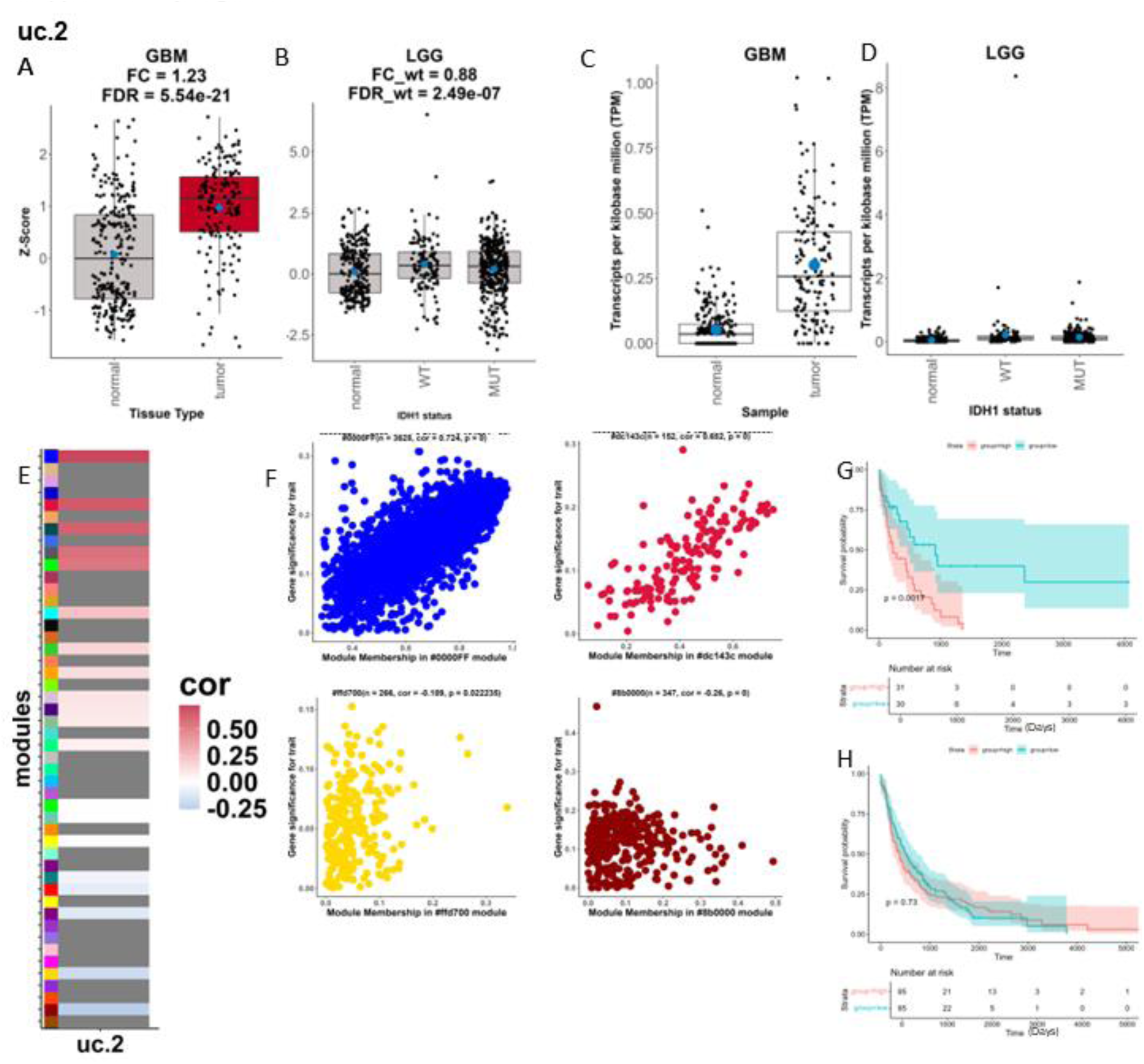
An exploration of a putative oncogenic TUCR, uc.2 in gliomas. A) Box- and dotplot showing uc.2 deregulation in GBM and B) LGG. Facets represent disease type. Red boxes represent upregulated TUCRs. Gray boxes represent TUCRs that are not deregulated. C) Box- and dotplot showing uc.2 absolute expression in GBM and D) LGG. Facets represent disease type. D) Heatmap depicting uc.2 gene module association. Positive correlations are red, while negative correlations are blue, with weak correlations in white. Modules with no linkage are gray. F) Scatter plots depicting uc.2 association with top 3 positive (top row) and negative (bottom row) correlation modules. G) Kaplan-Meier showing uc.2 association with GBM prognosis. Red line represents the TUCR high expression group. Teal line represents the TUCR low expression group. H) Kaplan-Meier showing uc.2 association with LGG prognosis. Red line represents the TUCR high expression group. Teal line represents the TUCR low expression group. (Similar figures for other putative oncogenic TUCRs available at in the TUCR Database provided as supplementary materials)

**Supplementary Figure 7.**
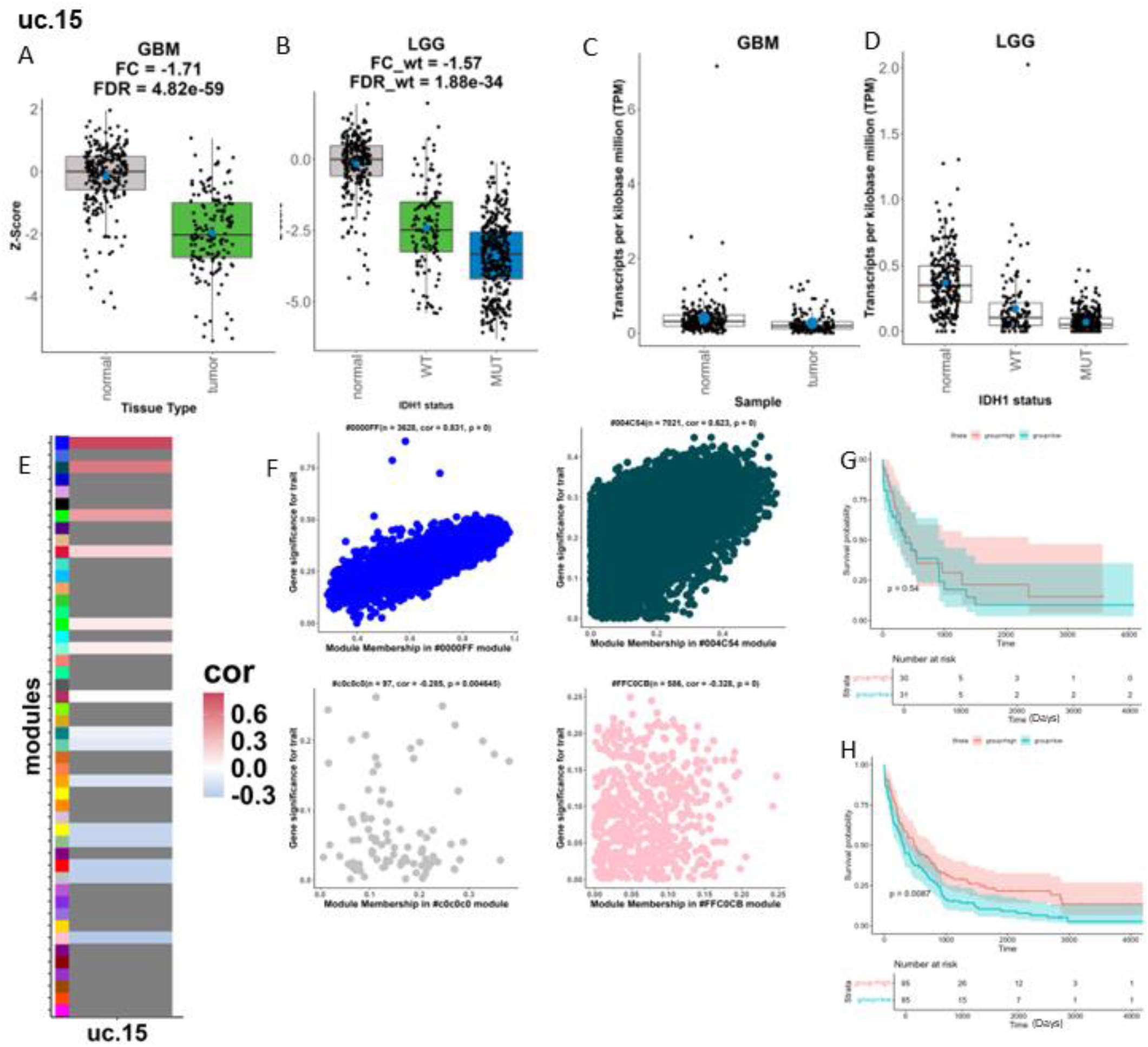
An exploration of a putative oncogenic TUCR, uc.15 in gliomas. A) Box- and dotplot showing uc.15 deregulation in GBM and B) LGG. Facets represent disease type. Red boxes represent upregulated TUCRs. Green boxes represent downregulated TUCRs. Gray boxes represent TUCRs that are not deregulated. C) Box- and dotplot showing uc.15 absolute expression in GBM and D) LGG. Facets represent disease type. E) Heatmap depicting uc.15 gene module association. Positive correlations are red, while negative correlations are blue, with weak correlations in white. Modules with no linkage are gray. F) Scatter plots depicting uc.15 association with top 3 positive (top row) and negative (bottom row) correlation modules. G) Kaplan-Meier showing uc.15 association with GBM prognosis. Red line represents the TUCR high expression group. Teal line represents the TUCR low expression group. H) Kaplan-Meier showing uc.15 association with LGG prognosis. Red line represents the TUCR high expression group. Teal line represents the TUCR low expression group. (Similar figures for other putative oncogenic TUCRs available at in the TUCR Database provided as supplementary materials)

**Supplementary Figure 8.**
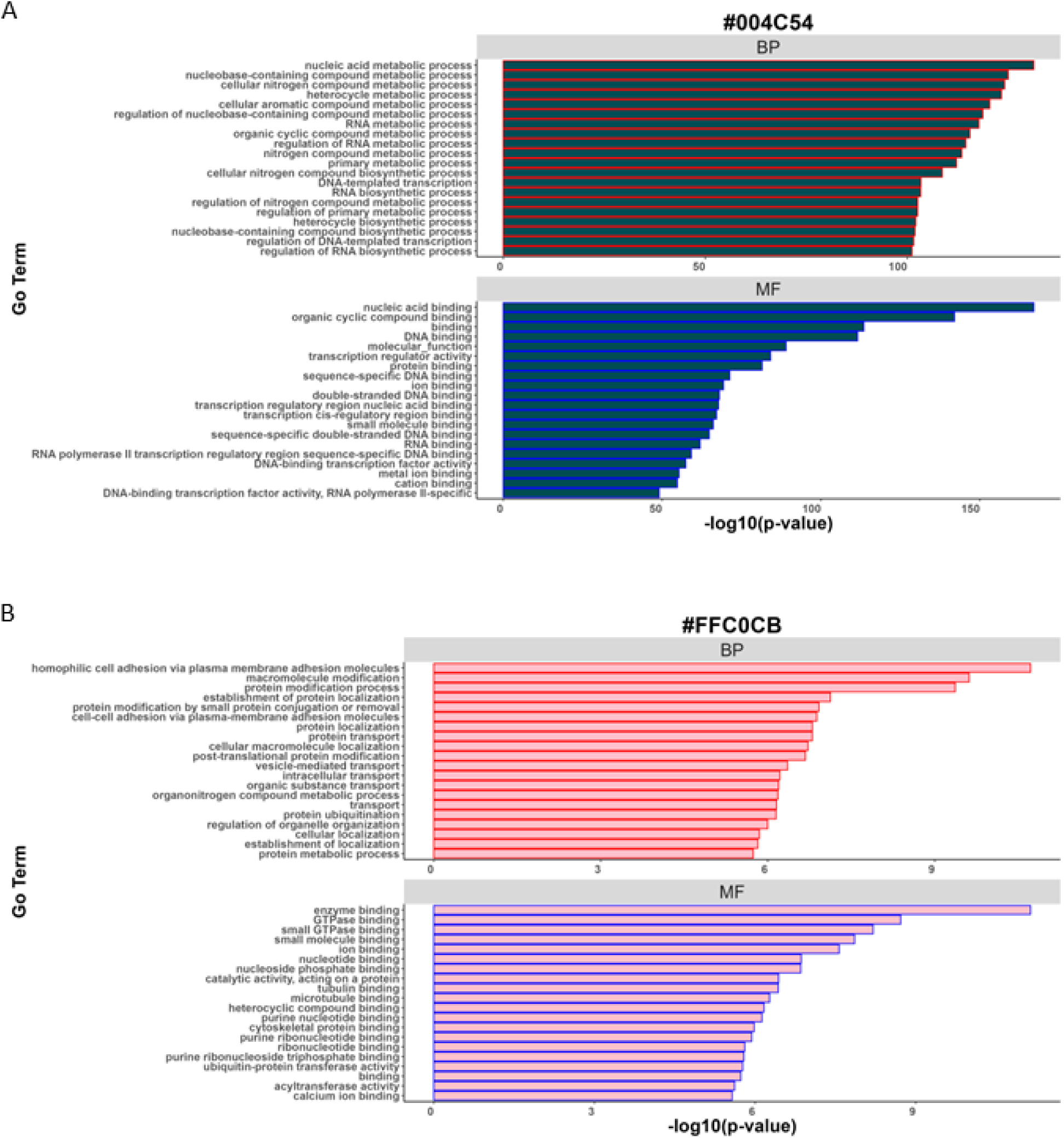
Top positively correlated TUCR modules in gliomas. A) The #004C54 module (midnight green) module is the most positively correlated with TUCRs. A) The is #FFC0CB (pink) the second most positively correlated module with TUCRs. (Hex codes for all modules are provided as supplement “colorcodelist”.

**Supplementary Figure 9.**
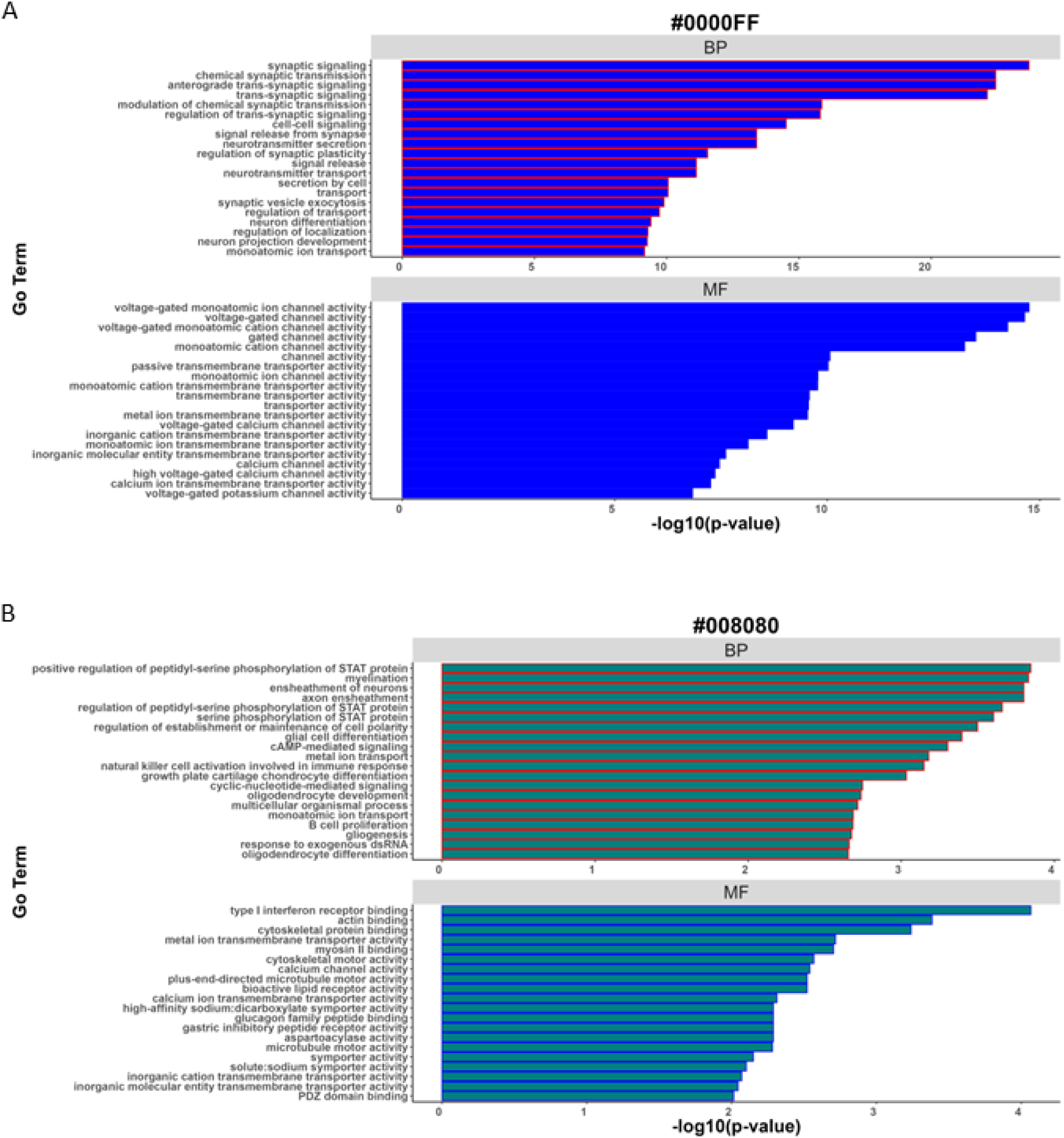
Top positively correlated TUCR modules in gliomas. A) The #f4a460 (sandybrown) module is the most positively correlated with TUCRs. B) The #FFA500 module (orange) is the second most positively correlated module with TUCRs. Gene ontology terms for all modules are provided as supplement. (TUCR Database)

**Supplementary Figure 10.**
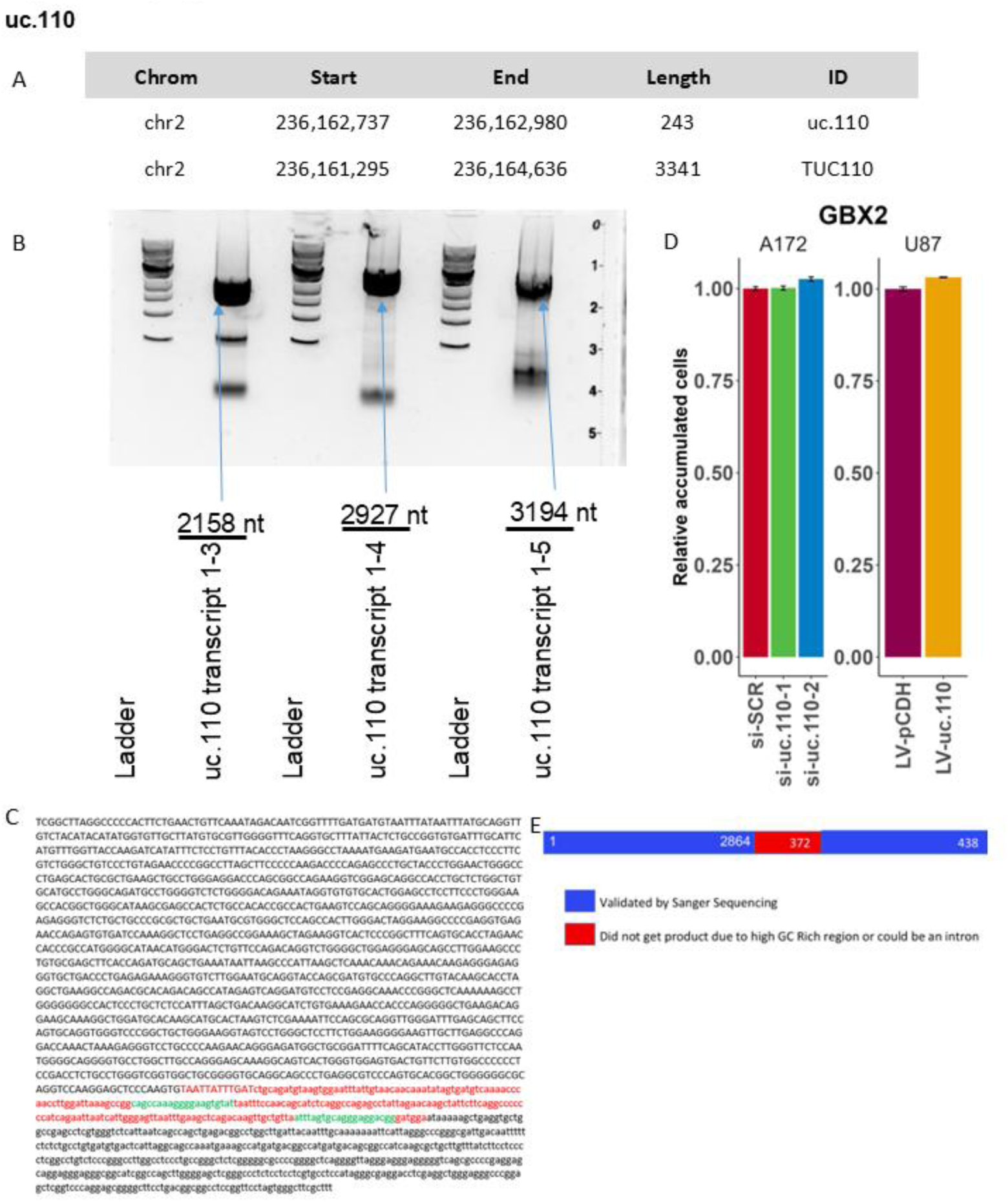
The uc.110 TUCR is part of a larger ncRNA transcript. A) We used de novo transcript reassembly of TCGA glioma RNA-Seq data and experimental PCR validation to identify the predicted sequence for the novel full RNA transcript containing uc.110. Table depicts uc.110 ultraconserved and predicted full transcript genomic locations and length in nucleotides (nt). B) PCR gel electrophoresis demonstrating validated uc.110 transcript variants. C) The validated full sequence of the 2,158 nt uc.110 transcript is provided. The ultraconserved uc.110 region is colored red and primer sequences are colored green. D) Bar chart confirmed uc.110 transcript independence from “host gene” GBX2 via qPCR. E) Cartoon demonstrating TUCR transcript independence GBX2 transcript.

**Supplementary Figure 11.**
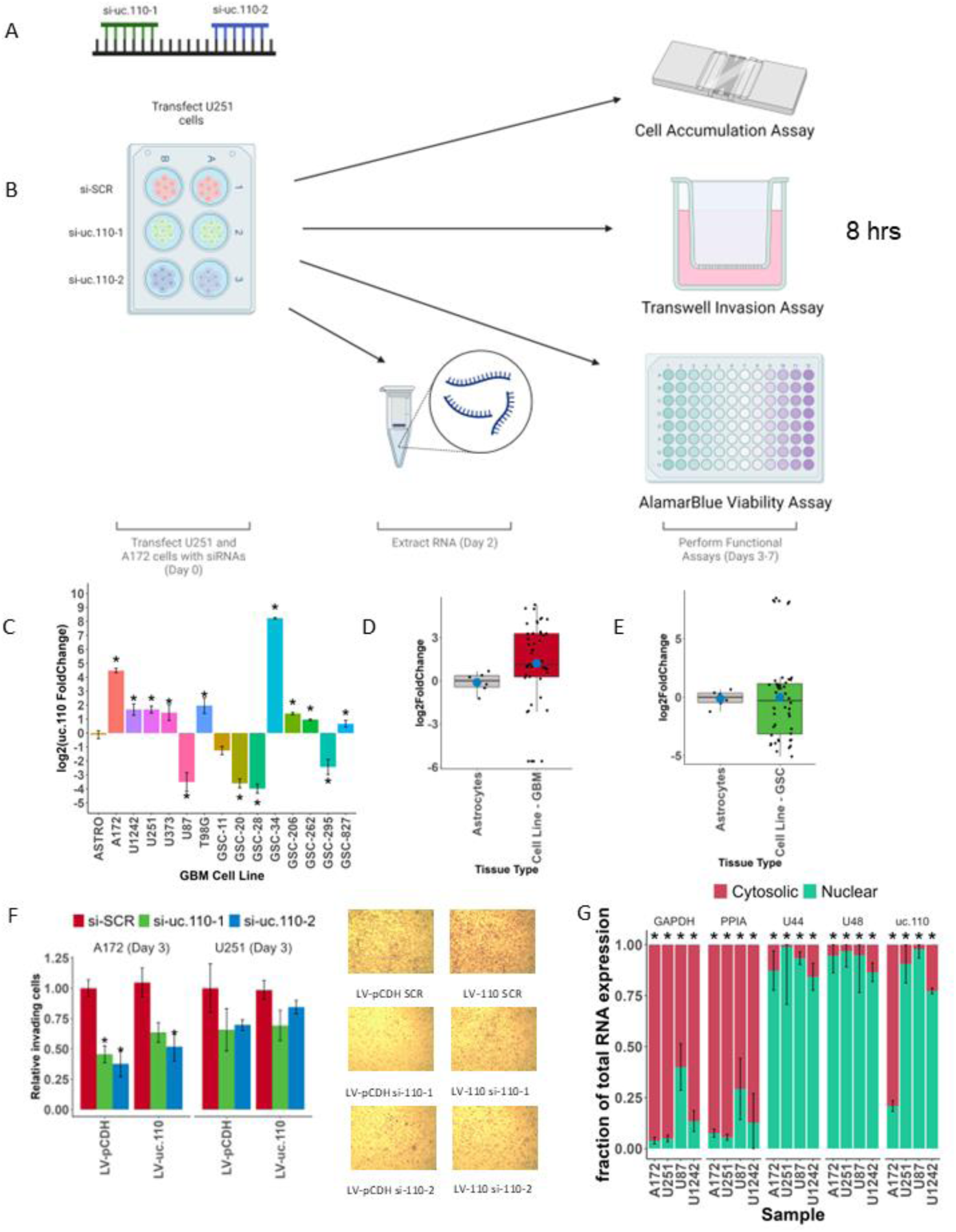
The uc.110 TUCR operates as an oncogene (cont.) A) Cartoon depicting two siRNAs that target different regions of the uc.110 TUCR. One starts at nt 96/243 (blue), and the other at nt 195/243 (green). B) Cartoon schematic depicting transfection protocol. Cells were transfected with siRNAs using Lipofectamine 2000 at D0. RNA was collected at D2, and functional assays were performed from D3-D7. C) Bar graph depicting uc.110 upregulation in banked GBM cell lines. D) Boxplot representing uc.110 expression in pooled glioma adherent cell lines versus normal human astrocytes. Red boxes indicate an upregulated TUCR. D) Boxplot representing uc.110 expression in pooled glioma adherent cell lines versus normal human astrocytes. Red boxes indicate an upregulated TUCR. E) Boxplot representing uc.110 expression in pooled glioma stem cell lines versus normal human astrocytes. Green boxes indicate a downregulated TUCR. F) Bar graph depicting that the cell invasion phenotype is rescued in A172 and U251 cells with uc.110 overexpression in the presence of siRNAs. Images are representative of the listed sample. si-SCR = scrambled control siRNA (red), si-uc.110-1 = siRNA targeting uc.110 at nucleotide 96/243 (green), si-uc.110-2 = siRNA targeting uc.110 at nucleotide 195/243 (blue). G) Cell fractionation bar graph depicting that uc.110 is a predominantly nuclear RNA molecule, with some cytosolic expression in A172s cells. Facets represent cytosolic (red) control genes (GAPDH,PPIA), nuclear (teal) control genes (U44,U48), and the uc.110 TUCR. * = p < 0.05

**Supplementary Figure 12.**
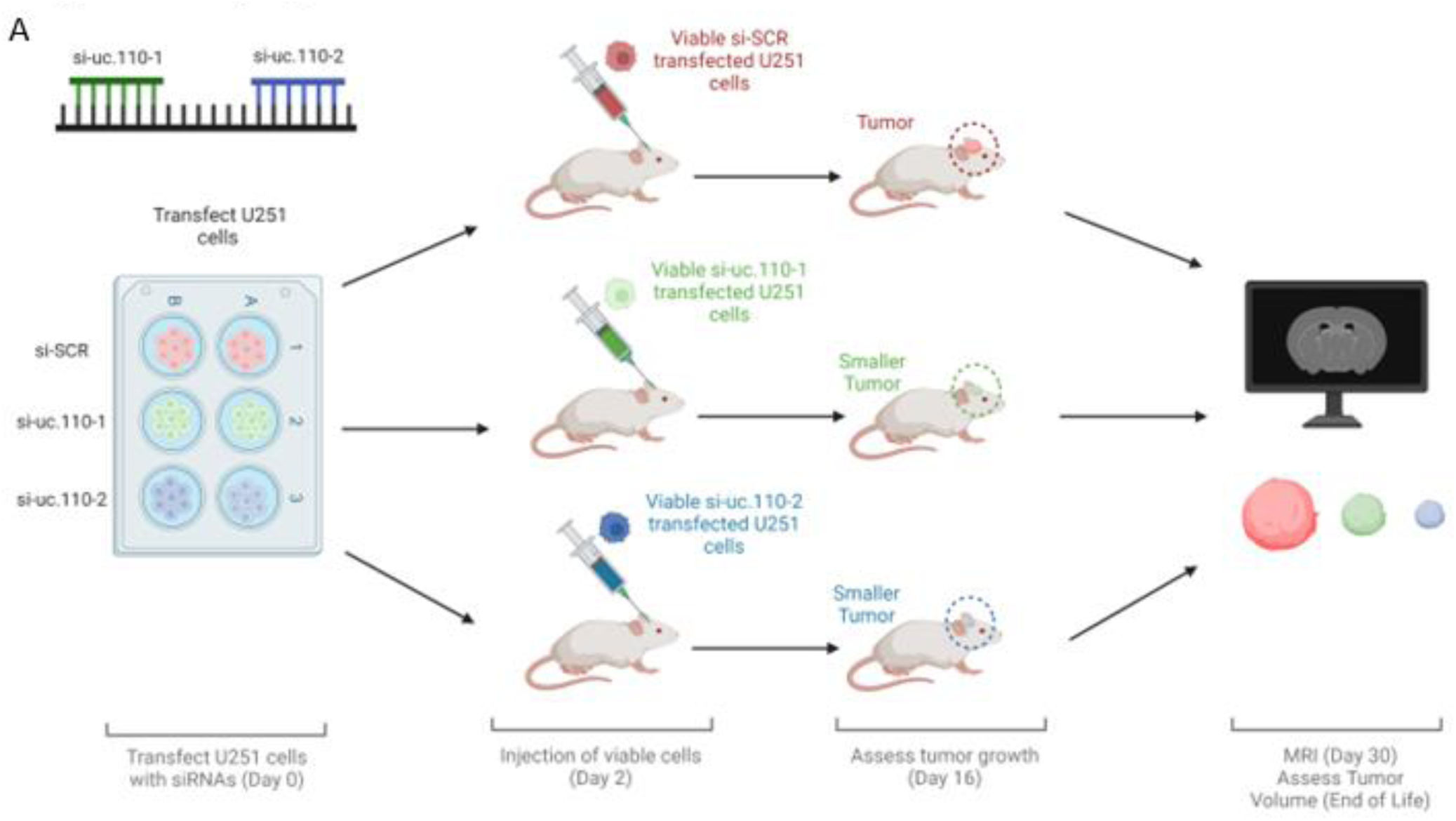
The uc.110 TUCR promotes tumor growth in vivo. A) Cartoon depiction of mouse experiment workflow. Cells were transfected with siRNAs using Lipofectamine 2000 at D0 and injected into mice at D2. Tumor growth was assessed weekly, starting at D16, via MRI.

**Supplementary Figure 13.**
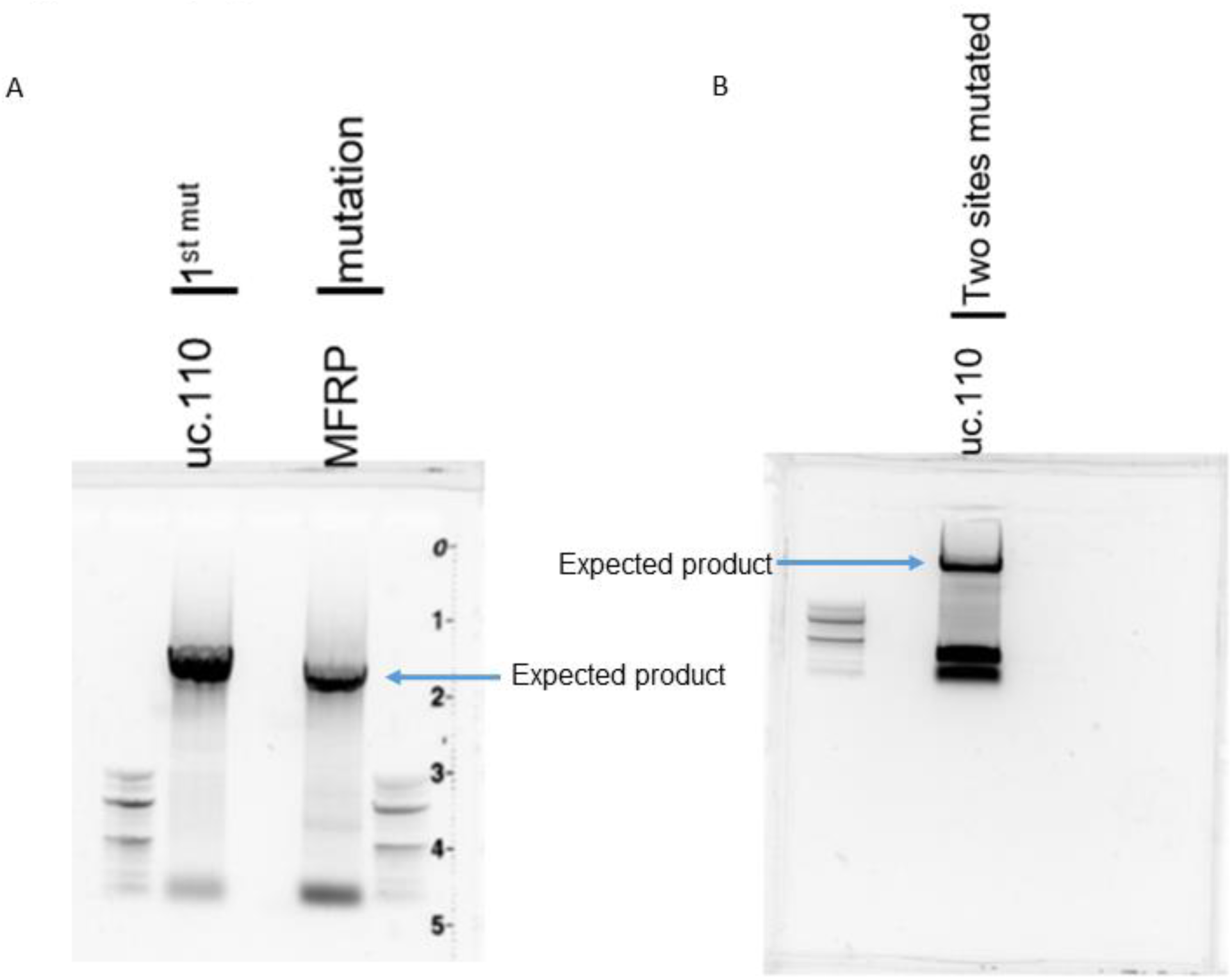
PCR confirmation of mutation of miR-544 binding sites for MFRP and uc.110. A) PCR gel showing expected products from uc.110 (first site) and MFRP mutations. B) PCR gel showing expected product from the second miR-544 binding site in uc.110

**Supplementary Figure 14.**
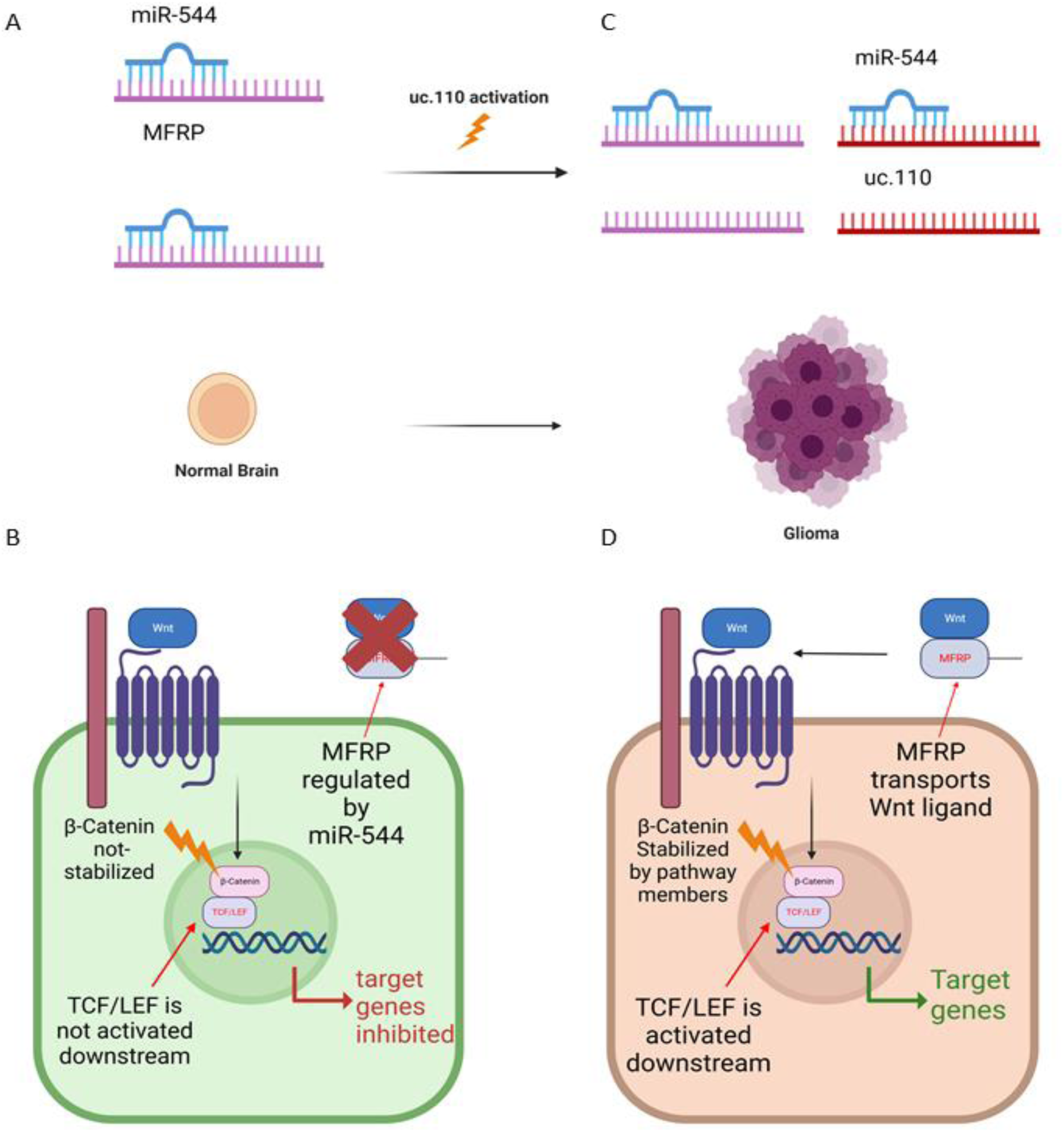
The uc.110 TUCR activates Wnt-signaling by sponging miR-544 from membrane frizzled related protein (MFRP) 3’UTR. A) Schematic depicting model for miR-544 sponging by uc.110. B) Schematic depicting simplified repressed Wnt-signaling pathway. In the normal brain, MFRP is downregulated by miR-544 as depicted in 13A. C) Activation of uc.110 in glioma tumors leads to decreased bioavailability of miR-544. This increases the bioavailability of MFRP. D) Schematic depicting simplified activated Wnt-signaling pathway. When MFRP bioavailability is increased by uc.110 activation, as depicted in 14C, Wnt-signaling is also increased.

## MATERIALS AND METHODS

### Troubleshooting

Please refer to README.md document on github.com/abounaderlab/tucr_project for additional instructions for downloading and extracting repositories, code, and supplementary data, and the corresponding author for any data access questions.

### Data Availability Statement

RNA-Seq data for Figure 5A will be made available on the Gene Expression Omnibus (GEO) prior to publication. Detailed methodologies and processed data is provided as supplement to this document (codebook.zip) and online at github.com/abounaderlab/tucr_project. Detailed TUCR results can be found in supplementary materials (TUCR Database), can be fully regenerated using the provided supplemental codebook (codebook.zip), and can also be found online at www.abounaderlab.org/.

Please refer to README.md document on github.com/abounaderlab/tucr_project for additional instructions for downloading and extracting repositories, code, and supplementary data, and the corresponding author for any data access questions.

### Detailed Computational Methodologies - Codebook

**Detailed methods, including access to information for all datasets used, can be found as a codebook provided in the supplementary materials and online at: github.com/abounaderlab/tucr_project.** This repository contains the processed data used to generate each figure and the codes to regenerate each figure in the manuscript, with a Markdown document for each figure with section headers. The UNIX command line (bash shell) and/or R programming languages were used for all analyses.

Please refer to README.md document on github.com/abounaderlab/tucr_project for additional instructions for downloading and extracting repositories, code, and supplementary data, and the corresponding author for any data access questions.

### TUCR Annotations [29, 30]

TUCR annotations were performed manually by overlaying consensus TUCR genomic annotation tracks to the hg38 human genome in the UCSC Genome Browser. TUCRs that were outside of an annotated gene were manually annotated as intergenic, while TUCRs that were within visible exons and introns were labeled as such. TUCRs that spanned multiple exons and introns were labeled as exonic/intronic. This was repeated for all 481 TUCRs.

In parallel, the bedtools closest package was used to identify TUCR host genes and TUCRs that are intergenic or intragenic. TUCRs with a distance of 0 nt from the nearest annotated genomic element were labeled as intragenic, while any TUCRs with a distance >0 were labeled intergenic. TUCRs These results were then cross referenced with the first analysis using table joining in R and manual confirmation to identify a consensus genomic annotation for each TUCR.

R/Rstudio was used for all downstream analyses. Detailed methods can be found as a codebook provided in the supplementary materials and online at: **github.com/abounaderlab/tucr_project**

### TUCR Chromatin Landscaping

U87 H3K4me3, RNA Pol.II, and H3K27ac CHIP-Seq data and U87 ATAC-Seq data were acquired from the Gene Expression Omnibus (GSE36354). SRA files were extracted via the SRA Toolkit, converted to fastq using fastq-dump, aligned to the hg38 genome via bowtie2, converted to sam using samtools, and converted to BAM format using bedtools. Features were extracted using macs2. Randomized control TUCRs were generated using Quinlan Labs’ bedtools [31, 32] and performing the shuffle command on the hg38 TUCR genomic positions, shuffling them throughout the genome randomly while retaining their same gene lengths. [31, 32] Bedtools fisher and R/RStudio [53, 54] were used to perform chi-square tests to compare predicted overlaps of actual peaks to expected peaks. R/Rstudio was used for all downstream analyses.

Detailed methods can be found as a codebook provided in the supplementary materials and online at: github.com/abounaderlab/tucr_project

### TCGA AND GTEx RNA-Seq Data [33, 34]

GBM (n = 161) and LGG (n = 504) RNA-Seq data were acquired from the Cancer Genome Atlas via the dbGAP portal and were compared to normal brain cortex from TCGA and the Genotype-Tissue Expression Database (GTEx, n = 260) acqu8ired via the ANVIL portal. Bowtie2 was used to align raw reads to the hg38 genome. Bedtools were used to convert files to sam, bam, and bed formats. R/Rstudio was used for all downstream analyses. LGG samples were subset into separate cohorts, by IDH1 mutant (n = 122) or wildtype (n = 382) status, for all downstream analyses, and all IDH1 mutant GBM samples (n = 8) were removed from the GBM cohort.

Detailed methods can be found as a codebook provided in the supplementary materials and online at: github.com/abounaderlab/tucr_project

### TUCR Expression, Deregulation, and Survival Analyses [33, 34, 54, 55]

TUCR expression, deregulation, and survival analyses, were analyzed using processed TCGA and GTEx RNA-Seq data and a workflow using R/RStudio. IDH1 mutant and wildtype samples were separated in different cohorts, and IDH1 GBM samples were removed from the GBM cohort, for these analyses. TPM scores were generated to assess absolute expression. DESeq2 and z-scores were used to calculate deregulated TUCRs. The cox coefficient of proportional hazards (CPH)from survminer Kaplan-Meier charts were used to assess correlation with patient outcomes.

Detailed methods can be found as a codebook provided in the supplementary materials and online at: github.com/abounaderlab/tucr_project

### TUCR weighted gene correlation network analysis (WGCNA) [36]

TUCR WGCNA was performed using processed TCGA and GTEx RNA-Seq data using a modified version of the R/RStudio workflow designed by Drs. Peter Langfelder and Steve Horvath at UC Los Angeles. TUCR expression data were converted to “trait data” for the workflow using TCGA GBM samples (n=166) compared to TCGA and GTex normal brain samples (n = 260). TUCR gene correlation modules were created by correlating TUCR expression as “trait data” to 42,644 annotated genomic elements. TUCR functions were predicted by identifying the most highly correlated modules with each TUCR and determining the functions of correlated genes within that module, as these genes are likely to have similar functions to the TUCR.

Detailed methods can be found as a codebook provided in the supplementary materials and online at: github.com/abounaderlab/tucr_project

### De novo transcript reassembly and validation [35]

De novo transcript assembly was performed on processed TCGA GBM and LGG RNA-Seq data using previously outlined protocols. The resultant bam files were then passed through the *stringtie* bioinformatics package to identify all transcripts in a reference file independent manner. TUCRs were intersected with these results using bedtools intersect. Intersections for intergenic transcripts were considered novel. For the transcript derived for uc.110 from this analysis, the transcript sequence was experimentally validated using sequential using PCR:

10 min at 95°C, followed by 40 cycles of 10 seconds at 95°C and 1 minute at 60°C. This PCR product was then outsourced for Sanger sequencing with GeneWiz from Azenta for the final sequenced product.

Detailed methods can be found as a codebook provided in the supplementary materials and online at: github.com/abounaderlab/tucr_project

### Patient Samples

GBM Tumor samples were acquired from the UVA Tumor Bank. Detailed patient information can be found as a supplement (Supplementary Table 2).

### Cell Lines and glioma stem cells

U87, U251, A172, and T98G glioblastoma cell lines were used in *in vitro* experiments and were acquired from ATCC. U87 cells were cultured in 500 mL minimum essential media (MEM) Earles (Gibco, #.11095-080) containing 5 mL penicillin/streptomycin (pen/strep, Gibco, Cat #.15140-133), 5 mL MEM non-essential amino acids (NEAA, Gibco, #.11140-050), 5 mL sodium pyruvate (Gibco, 100 nM, #.11360-070), 10 mL sodium bicarbonate (Gibco, 7.5%, #.25080-094), and 50 mL fetal bovine serum (FBS). T98G cells were cultured in 500 mL MEM Earles media containing 5 mL pen/strep, 5 mL NEAA, 5 mL sodium pyruvate, and 50 mL FBS. A172 cells were cultured in 500 mL Dulbecco’s modified eagle media (DMEM, Gibco, #.11965-092) containing 5 mL pen/strep, and 50 mL FBS. U251 cells were cultured in 500 mL RPMI L-Glutamine media (Gibco, #.11875093) containing 5 mL pen/strep and 25 mL FBS. GSC-34 and GSC-28 glioblastoma stem cells were cultured in neurobasal (L-glutamine negative) media (Gibco, #.21103-049) containing 5 mL pen/strep, 5 mL B-27 (without Vit-A, Gibco, #.12587-010), 2.5 mL N-2 (Gibco, #.17502-048), 1 mL EGF, 1 mL FGF, and 1.25 mL L-Glutamine. All cell media contained in 5 µL Plasmocure reagent to prevent mycoplasma contamination.

### Primer and Oligo Design

Primers and siRNAs were designed using the Primer3 and Thermofisher design portals, respectively. uc.110 forward primer sequence is 5’-CAGCCAAAGGGGAAGTGTAT-3’, and the reverse sequence is 5’-CCGTCCTCCCTGCACTAAAT-3’.

MFRP forward primer sequence is 5’-GCATCTATTCATGTGGCAGGC-3’, and the reverse sequence is 5’-TACTCCGGACCCTCCAGTTG-3’.

The miR-544 precursor was ordered from Invitrogen (#.AM17100). Negative control oligos were ordered from Ambion (#.AM4635).

### uc.110 stable overexpression

The full uc.110 transcript from “de novo transcript reassembly and validation” was cloned into the pCDH-EF1-MCS-BGH-PGK-GFP-T2A-Puro vector (# CD550A-1) using stbl3 competent *e.coli* cells and ampicillin selection. Amplified vector was extracted using the miniprep kit (Qiagen, #.27106). 0.75 µg of this vector, 0.75 µg of psPAX2 lentiviral gag-pol packaging vector, and 0.5 µg of pMD.2G VSV-G enveloping protein was transfected in 6 µL X-tremeGENE transfection reagent (#.06366236001) into 293T cells per manufacturer instructions to generate a lentivirus that was transduced to U87, U251, and A172 cells in media without antibiotics. These cells were subjected to antibody (puromycin) selection for uc.110-positive cells at D3.

### uc.110 quantitative (q)PCR

Total RNA was isolated using the RNEasy+ kit (Qiagen, #.74134) according to manufacturer instructions. RNA concentration and purity was measured via nanodrop. 800 ng of cDNA was synthesized (BIORAD T100 Thermal Cycler) using the iScript (BIORAD, #. 1708890) synthesis kit per manufacturer instructions. A 20 µL reaction mixture was then created for each condition with the following concentrations: 1 µL of combined forward/reverse primers (5 µM), 10 µL of iQ SYBR Green master mix (#1798880), 4 µL of nuclease free water, and 5 µL of synthesized cDNA. These reactions were cycled (BIORAD CFX Real Time System) in 96-well plates: 10 min at 95°C, followed by 40 cycles of 10 seconds at 95°C and 1 minute at 60°C.

### Cell Counting (Accumulation) Assay [37–39, 44]

Cells were seeded in 6-well culture plates with full serum media at 30,000/well density at D-1. At D0, each well was transfected via master mix 3 µL of siRNAs (20 µM) via 9 µL Lipofectamine 2000 (Invitrogen, #.11668-019) in 300 µL OPTI-MEM (Gibco, #.31985-070) and 700 µL antibiotic and empty media for 6 hours. At 6 hours, media were replaced with fresh media containing antibiotics and FBS. Cells were then counted via haemocytometer at Days 1, 3, 5, and 7 for each cell line.

### Transwell Cell Invasion/Migration Assay [42–44]

Cells were seeded in 6-well culture plates with full serum media at 300k/well density at D-1. At D0, each well was transfected via master mix 3 µL of siRNAs (20 µM) via 9 µL Lipofectamine 2000 in 300 µL OPTI-MEM and 700 µL antibiotic and empty media for 6 hours. At 6 hours, the media were replaced with fresh media containing antibiotics and FBS. The cells were then seeded in empty media at 200k/chamber into Transwell Invasion Chambers coated with Collagen IV.

After 8 hours, non-invading cells were cleared and invading cells were stained with Crystal Violet.

### Alamar Blue Cell Viability Assay [40–41]

Cells were seeded in 96-well culture plates with full serum media at 10k/well density at D-1. Border wells were filled with media to account for edge effects. At D0, each well was transfected via master mix 1 µL of siRNAs (20 µM) via 3 µL Lipofectamine 2000 in 30 µL OPTI-MEM and 70 µL antibiotic and empty media for 6 hours. At 6 hours, media were replaced with fresh media containing antibiotics and FBS. Functional assays were performed using the AlamarBlue kit (Life Technologies #. A50100) per manufacturer instructions. Reactions were allowed to proceed for 1 hour.

### Ex vivo knockdown of uc.110

Cells were seeded in 6-well culture plates with full serum media at 300k/well density at D-1. At D0, each well was transfected with 3 µL of siRNAs (20 µM) via 9 µL Lipofectamine 2000 in 300 µL OPTI-MEM and 700 µL antibiotic and empty media for 6 hours. At 6 hours, the media were replaced with fresh media containing antibiotics and FBS. Mouse experiments were performed using xenograft models and intracranial injections of U251 cells post transfection with siRNA oligonucleotides. Cells were injected at D2 and were imaged at two-week intervals via MRI.

Survival was assessed daily and tumor volume was measured at the end of life.

### Characterization of transcriptome post-uc.110 knockdown

Cells were seeded in 6-well culture plates with full serum media at 300k/well density at D1. At D0, each well was transfected with 3 µL of siRNAs (20 µM) using 9 µL Lipofectamine 2000 in 300 µL OPTI-MEM and 700 µL antibiotic and empty media for 6 hours. At 6 hours, the media were replaced with fresh media containing antibiotics and FBS. RNA Libraries were collected and sequenced via RNA-Seq on Day 2 (post transfection).

### Luciferase Reporter Vector Construction

The Luciferase reporter vector were constructed via insertion of uc.110 conserved region and 3’UTR of MFRP downstream of Renilla luciferase stop codon in psi-CHECK2 dual luciferase vectors (Promega, Madison, WI, USA). The insertions were validated by sequencing. Uc.110 and MFRP primer pairs with XholI and NotI sequence at 5’ and 3’ respectively, uc.110-FW: 5’-ATATATctcgagCGAGGTGAGAACCAGAGTGT-3’, uc.110-RW: 5’-AATAATgcggccgcTTGGCTGCCTAATGAGTCACA-3’, MFRP-FW: 5’-ATATATctcgagAAATGGGGTCTGGTCCTTGG-3’ and MFRP-RW: 5’-AATAATgcggccgcTCGCCTTTCTCTCCCGGA-3’ were used for PCR amplification. Site-directed mutagenesis of predicted miR-544 target sites for both uc.110 and MFRP were performed to generate mutant vectors.

### 3’UTR Reporter Assays

To determine whether miR-544 directly binds to the MFRP 3’UTR and uc.110, cells were transfected with miR-544 or miR-scr (control) for 24 hour. The cells were then transfected with luciferase reporter control or 3’UTR-MFRP or uc.110 as well as corresponsive mutant vectors for 24 hours. Luciferase assays wered performed using the Luciferase System Kit (Promega) and luminescence was measured. Renilla luciferase activity was double normalized by dividing each well first by firefly activity and then by average luciferase/firefly value in a parallel set done with constitutive luciferase plasmid.

### TCF/LEF reporter Assays [57–58]

Cells were seeded in 6-well culture plates with full serum media at 300k/well density at D-1. At D0, each well was transfected with 3 µL of siRNA/miRNA (20 µM) using 9 µL Lipofectamine 2000 in 300 µL OPTI-MEM and 700 µL antibiotic and empty media for 6 hours. At 6 hours, the media were replaced with fresh media containing antibiotics and FBS. MFRP and uc.110 sequences were cloned into the PROMEGA pmirGLO Luciferase vector (E1330). BPS Dual reporter luciferase assays were ordered for TCF/LEF (#.60500) and uc.110/MFRP (#.60683) experiments.

### In Vivo Tumor Formation

Adult male and female Nude: Hsd:Athymic Nude-Foxn1 mice were purchased from Harlan. All the animal work was conducted at the Animal Research Core Facility at the University of Virginia School of Medicine in accordance with the institutional guidelines. Mice used for this study were anesthetized with ketamine (17.4 mg/20g), xylazine (2.6 mg/20g) and placed on a sterotactic frame. Tumor xenografts were generated by implantation of U251 cells transfected with si-uc.110-1, si-cu.110-2 or si-Scr. U251 cells (3×10^5^ cells; n=5) were stereotactically implanted into mice in their right striata at the coordinates from the bregma 1mm anterior, 1.5 mm lateral and 2.5 mm intraparenchymal. Three weeks after tumor implantation, the animals were subjected to brain MRI. To measure the tumor size, 20 ul of gadopentetate dimeglumine (Magnevist, Bayer Healthcare) was intraperitoneally injected 15 minutes before scanning. Tumor volumes were measured using MicroDicom.

### Statistical Analyses

Comparisons between means of samples were performed using Student’s t-test and one-way ANOVAs. Comparisons between categorical variables were performed using chi-squared and Fisher’s exact test. Comparisons were considered statistically significant if the p-value was less than 0.05. Multiple hypothesis correction using the Bonferroni method was performed for all comparisons, converting raw p-values to FDR, unless otherwise stated. Molecular experiment tests were performed in SigmaPlot 14.0, while computational experiment tests were performed using bedtools and/or RStudio.

Detailed methods can be found as a codebook provided in the supplementary materials and online at: github.com/abounaderlab/tucr_project

## AUTHOR CONTRIBUTIONS

Author contributions are defined using Elsevier’s CRediT format:

⮚ **Myron Gibert Jr:** Conceptualization, Methodology, Software, Validation, Formal Analysis,
⮚ Investigation, Writing, Visualization, Supervision, Project Administration, Funding Acquisition
⮚ **Ying Zhang:** Methodology, Investigation, Resources, Formal Analysis, Data Curation, Review
⮚ and Editing
⮚ **Shekhar Saha:** Methodology, Investigation, Resources, Formal Analysis, Data Curation, Review
⮚ and Editing
⮚ **Guruprasad Varma Konduru:** Methodology, Software, Validation, Formal Analysis, Resources, Data Curation
⮚ **Pawel Marcinkiewicz:** Investigation, Resources, Formal Analysis, Data Curation, Review and
⮚ Editing
⮚ **Sylwia Bednarek:** Methodology, Validation, Formal Analysis, Investigation, Review and Editing,
⮚ Visualization
⮚ **Collin Dube:** Methodology, Writing, Visualization, Review and Editing
⮚ **Kadie Hudson:** Methodology, Writing, Visualization, Review and Editing
⮚ **Yunan Sun:** Methodology, Writing, Visualization, Review and Editing
⮚ **Bilhan Chagari:** Formal Analysis, Investigation, Writing, Data Curation
⮚ **Aditya Sarkar:** Formal Analysis, Investigation, Writing, Data Curation
⮚ **Christian Roig-Laboy:** Formal Analysis, Investigation, Writing, Data Curation
⮚ **Natalie Neace:** Formal Analysis, Investigation, Writing, Data Curation
⮚ **Karim Saoud:** Formal Analysis, Investigation, Writing, Data Curation
⮚ **Initha Setiady:** Formal Analysis, Investigation, Writing, Data Curation
⮚ **Farina Hanif:** Methodology, Investigation, Review and Editing
⮚ **David Schiff:** Tumor tissue contribution, Review and Editing
⮚ **Benjamin Kefas:** Methodology, Investigation, Review and Editing
⮚ **Markus Hafner:** Conceptualization and editing
⮚ **Pankaj Kumar:** Methodology, Software, Validation, Formal Analysis, Resources, Data Curation
⮚ **Roger Abounader:** Conceptualization, Methodology, Resources, Review and Editing, Supervision, Project Administration, Funding Acquisition

## ACKNOWLEDGMENTS

This article was supported by NIH UO1 CA220841, NINDS 1R21NS122136-01, NINDS RO1 NS122222, NCI Cancer Center Support Grant 5P30CA044579, and a Schiff Foundation grant (all to R.A.). The work was also supported by the UVA Cancer Center Bioinformatics Core and the Molecular Imaging Core. We thank the dbGAP and the TCGA data management teams for providing access to raw RNAseq data. We would also like to express our gratitude to the patients for their participation in TCGA and the UVA tumor repository. Special thanks goes to Drs. Stephen Turner and Alex Koeppel for their work with the Bioinformatics Core and early contributions to some of the computational studies.

## Notes

https://github.com/abounaderlab/tucr_project

### Competing Interest Statement

The authors have declared no competing interest.

### Summary of Updates

Manuscript updated to reflect most recent developments/findings.

## REFERENCES

1. Bejerano, G., et al., Ultraconserved elements in the human genome. Science, 2004. 304(5675): p. 1321–5.

2. Gibert, M.K., Jr., et al., Transcribed Ultraconserved Regions in Cancer. Cells, 2022. 11(10).

3. Calin, G.A. and C.M. Croce, Chronic lymphocytic leukemia: interplay between noncoding RNAs and protein-coding genes. Blood, 2009. 114(23): p. 4761–70.

4. Calin, G.A., et al., Ultraconserved regions encoding ncRNAs are altered in human leukemias and carcinomas. Cancer Cell, 2007. 12(3): p. 215–29.

5. Colwell, M., et al., Evolutionary conservation of DNA methylation in CpG sites within ultraconserved noncoding elements. Epigenetics, 2018. 13(1): p. 49–60.

6. Edwards, J.K., et al., MicroRNAs and ultraconserved genes as diagnostic markers and therapeutic targets in cancer and cardiovascular diseases. J Cardiovasc Transl Res, 2010. 3(3): p. 271–9

7. Fabris, L. and G.A. Calin, Understanding the Genomic Ultraconservations: T-UCRs and Cancer. Int Rev Cell Mol Biol, 2017. 333: p. 159–172.

8. Lujambio, A., et al., CpG island hypermethylation-associated silencing of non-coding RNAs transcribed from ultraconserved regions in human cancer. Oncogene, 2010. 29(48):p. 6390–401.

9. McCole, R.B., et al., Abnormal dosage of ultraconserved elements is highly disfavored in healthy cells but not cancer cells. PLoS Genet, 2014. 10(10): p. e1004646.

10. Scaruffi, P., The transcribed-ultraconserved regions: a novel class of long noncoding RNAs involved in cancer susceptibility. ScientificWorldJournal, 2011. 11: p. 340–52.

11. Zambalde, E.P., et al., Transcribed Ultraconserved Regions Are Associated with Clinicopathological Features in Breast Cancer. Biomolecules, 2022. 12(2).

12. Braicu, C., et al., The Function of Non-Coding RNAs in Lung Cancer Tumorigenesis.Cancers (Basel), 2019. 11(5).

13. Catana, C.S., et al., Non-coding RNAs, the Trojan horse in two-way communication between tumor and stroma in colorectal and hepatocellular carcinoma. Oncotarget, 2017. 8(17): p. 29519–29534.

14. Irimie, A.I., et al., The Unforeseen Non-Coding RNAs in Head and Neck Cancer. Genes (Basel), 2018. 9(3).

15. Ling, H., et al., Junk DNA and the long non-coding RNA twist in cancer genetics. Oncogene, 2015. 34(39): p. 5003–11.

16. Liz, J., et al., Regulation of pri-miRNA processing by a long noncoding RNA transcribed from an ultraconserved region. Mol Cell, 2014. 55(1): p. 138–47.

17. Sakamoto, N., et al., Non-coding RNAs are promising targets for stem cell-based cancer therapy. Noncoding RNA Res, 2017. 2(2): p. 83–87.

18. Kiran, M., et al., A Prognostic Signature for Lower Grade Gliomas Based on Expression of Long Non-Coding RNAs. Mol Neurobiol, 2019. 56(7): p. 4786–4798.

19. Dey, B.K., A.C. Mueller, and A. Dutta, Long non-coding RNAs as emerging regulators of differentiation, development, and disease. Transcription, 2014. 5(4): p. e944014.

20. Cruickshanks, N., et al., Role and Therapeutic Targeting of the HGF/MET Pathway in Glioblastoma. Cancers (Basel), 2017. 9(7).

21. Davis, M.E., Glioblastoma: Overview of Disease and Treatment. Clin J Oncol Nurs, 2016. 20(5 Suppl): p. S2–8.

22. Lathia, J.D., et al., Cancer stem cells in glioblastoma. Genes Dev, 2015. 29(12): p. 1203–17

23. Kalkan, R., Glioblastoma Stem Cells as a New Therapeutic Target for Glioblastoma. Clin Med Insights Oncol, 2015. 9: p. 95–103.

24. Thakkar, J.P., et al., Epidemiologic and molecular prognostic review of glioblastoma. Cancer Epidemiol Biomarkers Prev, 2014. 23(10): p. 1985–96.

25. Mao, H., et al., Deregulated signaling pathways in glioblastoma multiforme: molecular mechanisms and therapeutic targets. Cancer Invest, 2012. 30(1): p. 48–56.

26. Altaner, C., Glioblastoma and stem cells. Neoplasma, 2008. 55(5): p. 369–74.

27. Mourad, P.D., et al., Why are systemic glioblastoma metastases rare? Systemic and cerebral growth of mouse glioblastoma. Surg Neurol, 2005. 63(6): p. 511–9; discussion

28. Zhang, Y., et al., The p53 Pathway in Glioblastoma. Cancers (Basel), 2018. 10(9).

29. Raney, B.J., et al., Track data hubs enable visualization of user-defined genome-wide annotations on the UCSC Genome Browser. Bioinformatics, 2014. 30(7): p. 1003–5.

30. Kent, W.J., et al., The human genome browser at UCSC. Genome Res, 2002. 12(6): p. 996–1006.

31. Quinlan, A.R., BEDTools: The Swiss-Army Tool for Genome Feature Analysis. Curr Protoc Bioinformatics, 2014. 47: p. 11 12 1–34.

32. Quinlan, A.R. and I.M. Hall, BEDTools: a flexible suite of utilities for comparing genomic features. Bioinformatics, 2010. 26(6): p. 841–2.

33. Cancer Genome Atlas Research, N., et al., The Cancer Genome Atlas Pan-Cancer analysis project. Nat Genet, 2013. 45(10): p. 1113–20.

34. Stanfill, A.G. and X. Cao, Enhancing Research Through the Use of the Genotype-Tissue Expression (GTEx) Database. Biol Res Nurs, 2021. 23(3): p. 533–540.

35. Pertea, M., et al., CHESS: a new human gene catalog curated from thousands of largescale RNA sequencing experiments reveals extensive transcriptional noise. Genome Biol, 2018. 19(1): p. 208.

36. Langfelder, P. and S. Horvath, WGCNA: an R package for weighted correlation network analysis. BMC Bioinformatics, 2008. 9: p. 559.

37. Hanif, F., et al., miR-3174 Is a New Tumor Suppressor MicroRNA That Inhibits Several Tumor-Promoting Genes in Glioblastoma. Int J Mol Sci, 2023. 24(11).

38. Mulcahy, E.Q.X., et al., MicroRNA 3928 Suppresses Glioblastoma through Downregulation of Several Oncogenes and Upregulation of p53. Int J Mol Sci, 2022. 23(7).

39. Cruickshanks, N., et al., Discovery and Therapeutic Exploitation of Mechanisms of Resistance to MET Inhibitors in Glioblastoma. Clin Cancer Res, 2019. 25(2): p. 663–673.

40. Kumar, P., A. Nagarajan, and P.D. Uchil, Analysis of Cell Viability by the alamarBlue Assay. Cold Spring Harb Protoc, 2018. 2018(6).

41. Voytik-Harbin, S.L., et al., Application and evaluation of the alamarBlue assay for cell growth and survival of fibroblasts. In Vitro Cell Dev Biol Anim, 1998. 34(3): p. 239–46.

42. Stoellinger, H.M. and A.R. Alexanian, Modifications to the Transwell Migration/Invasion Assay Method That Eases Assay Performance and Improves the Accuracy. Assay Drug Dev Technol, 2022. 20(2): p. 75–82.

43. Liu, X. and X. Wu, Utilizing Matrigel Transwell Invasion Assay to Detect and Enumerate Circulating Tumor Cells. Methods Mol Biol, 2017. 1634: p. 277–282.

44. Justus, C.R., et al., In vitro cell migration and invasion assays. J Vis Exp, 2014(88).

45. Guessous, F., et al., Cooperation between c-Met and Focal Adhesion Kinase Family Members in Medulloblastoma and Implications for Therapy. Mol Cancer Ther, 2012. 11(2): p. 288–97.

46. Sonoda, Y., et al., Akt pathway activation converts anaplastic astrocytoma to glioblastoma multiforme in a human astrocyte model of glioma. Cancer Res, 2001. 61(18): p. 6674–8.

47. Won, J., et al., Membrane frizzled-related protein is necessary for the normal development and maintenance of photoreceptor outer segments. Vis Neurosci, 2008. 25(4): p. 563–74.

48. Katoh, M., Molecular cloning and characterization of MFRP, a novel gene encoding amembrane-type Frizzled-related protein. Biochem Biophys Res Commun, 2001. 282(1): p. 116–23.

49. Slaby, O., R. Laga, and O. Sedlacek, Therapeutic targeting of non-coding RNAs in cancer. Biochem J, 2017. 474(24): p. 4219–4251.

50. Saleembhasha, A. and S. Mishra, Long non-coding RNAs as pan-cancer master gene regulators of associated protein-coding genes: a systems biology approach. PeerJ, 2019. 7: p. e6388.

51. Li, L., et al., HOX cluster-embedded antisense long non-coding RNAs in lung cancer.Cancer Lett, 2019. 450: p. 14–21.

52. Jiang, D., et al., Long Chain Non-Coding RNA (lncRNA) HOTAIR Knockdown Increases miR-454-3p to Suppress Gastric Cancer Growth by Targeting STAT3/Cyclin D1. Med Sci Monit, 2019. 25: p. 1537–1548.

53. Soler, M., et al., The transcribed ultraconserved region uc.160+ enhances processing and A-to-I editing of the miR-376 cluster: hypermethylation improves glioma prognosis. Mol Oncol, 2022. 16(3): p. 648–664.

54. Team, R., RStudio: Integrated Development for R. 2020.

55. Team., R.D.C. R: A Language and Environment for Statistical Computing. 2012; Available from: http://www.r-project.org/.

56. Mestdagh P., Fredlund E., Pattyn F., Rihani A., Van Maerken T., Vermeulen J., Kumps C., Menten B., De Preter K., Schramm A., et al. An integrative genomics screen uncovers ncRNA T-UCR functions in neuroblastoma tumours. Oncogene. 2010;29:3583–3592. doi: 10.1038/onc.2010.106.

57. Clevers H (2006) Wnt/beta-catenin signaling in development and disease. Cell 127(3):469–480.

58. Chen B et al. (2009) Small molecule-mediated disruption of Wnt-dependent signaling in tissue regeneration and cancer. Nature Chemical Biology 5(2):100–107.

59. Lee, Y., Lee, JK., Ahn, S. et al. WNT signaling in glioblastoma and therapeutic opportunities. Lab Invest 96, 137–150 (2016). 10.1038/labinvest.2015.140

60. Latour M, Her NG, Kesari S, Nurmemmedov E. WNT Signaling as a Therapeutic Target for Glioblastoma. Int J Mol Sci. 2021 Aug 5;22(16):8428. doi: 10.3390/ijms22168428. PMID: 34445128; PMCID: PMC8395085.

